# Allopatric origin of sympatric whitefish morphs with insights on the genetic basis of their reproductive isolation

**DOI:** 10.1101/2021.09.11.459905

**Authors:** Bohao Fang, Paolo Momigliano, Kimmo K. Kahilainen, Juha Merilä

## Abstract

The European whitefish (*Coregonus lavaretus*) species complex is a classic example of recent adaptive radiation. Here we examine a whitefish population introduced to northern Finnish Lake Tsahkal in late 1960’s, where three divergent morphs (*viz*. littoral, pelagic and profundal feeders) were found ten generations after. Using demographic modelling based on genomic data we show that whitefish morphs evolved during a phase of strict isolation, refuting a rapid symmetric speciation scenario. The lake is now an artificial hybrid zone between morphs originated in allopatry. Despite their current syntopy, clear genetic differentiation remains between two of the three morphs. Using admixture mapping three quantitative trait loci associated with gonad weight variation, a proxy for sexual maturity and spawning time, were identified. We suggest that ecological adaptations in spawning time evolved in allopatry are currently maintaining partial reproductive isolation in the absence of other barriers to gene flow.

## Introduction

The pace at which evolution and speciation occur, and the mechanisms determining their tempo, have intrigued evolutionary biologists for a long time ((Simpson 1944; Bush *et al*. 1977; Kornfield 1978)), and still continue today (Schluter 2000; Nosil 2012; Lescak *et al*. 2015; Momigliano *et al*. 2017; Salzburger 2018; Skúlason *et al*. 2019). Concerns over the organims’ ability to adapt to environmental changes (e.g. Gienapp *et al*. (2008); Merilä & Hendry (2014)), and the realisation that adaptation can be very rapid (Thomas *et al*. 2017; Matthews *et al*. 2018; Marques *et al*. 2019b), have sparked new interest towards the rates of evolution. Adaptation from standing genetic variation can be fast, especially if genetic diversity has been augmented by historical gene flow (Häkli *et al*. 2018; Salzburger 2018; Jacobs *et al*. 2019; Marques *et al*. 2019a). One example is the evolution of freshwater-adapted three-spined sticklebacks (*Gasterosteus aculeatus*) colonising ponds that were formed during uplift caused by the 1964 Great Alaska Earthquake (Lescak *et al*. 2015). Similarly rapid adaptation has been observed for sockeye salmon and Darwin’s finches when colonizing new environments (Hendry *et al*. 2000; Lamichhaney *et al*. 2018). In all these examples, broad ecological opportunities likely facilitated rapid phenotypic and genetic differentiation.

Coregonid fishes have undergone extensive recent adaptive radiations but intermittent gene flow among morphs and species is common (Østbye *et al*. 2005; Bernatchez *et al*. 2010; Hudson *et al*. 2011; Vonlanthen *et al*. 2012). European whitefish (*Coregonus lavaretus*) experienced frequent adaptive radiations throughout their distribution (Svärdson 1979; Østbye *et al*. 2005; Vonlanthen *et al*. 2012). Facing different ecological opportunities, *C. lavaretus* evolved partially reproductively isolated, co-existing morphs utilizing different niches (Kahilainen & Østbye 2006; Siwertsson *et al*. 2010; Præbel *et al*. 2013). These morphs differ in several phenotypic traits (Harrod *et al*. 2010; Præbel *et al*. 2013), such as body size and number of gill rakers, a heritable traits related to feeding ecology (Kahilainen *et al*. 2011; Häkli *et al*. 2018). While rapid whitefish differentiation has been documented in both Nearctic and Palearctic regions (Bernatchez *et al*. 2010; Hudson *et al*. 2011), less is known about the genomic basis of their rapid phenotypic divergence (but see Vonlanthen *et al*. (2012); Jacobs *et al*. (2019); Rougeux *et al*. (2019a)).

In northern Finland, the Finnish Fisheries authorities facilitated introductions of whitefish in several high-altitude lakes including Lake Tsahkal, where the species did not previously occur, in 1960’s. Sampling in Lake Tsahkal in 2011 revealed three different morphs inhabiting littoral, pelagic and profundal habitats (Kahilainen *et al*. 2011; Præbel *et al*. 2013). There is evidence of both allopatric and sympatric speciation of whitefish morphs (Præbel *et al*. 2013), and genetic diversity fuelled by historical admixture can facilitate rapid phenotypic divergence when ecological opportunity arises (Jacobs *et al*. 2019). It is possible that the whitefish introduced to Lake Tsahkal came from a single ancestral population and gave rise to different morphs in ca. 50 years (≈10 generations). The alternative hypothesis is that the three morphs were simultaneously introduced in the lake. In such case, the whitefish of Lake Tsahkal would provide a unique opportunity to study the ecological and genetic bases of reproductive barriers following secondary contact.

Lake Tsahkal provides also an ideal system to study the genetic basis of phenotypic traits defining the whitefish morphs via admixture mapping (Gompert *et al*. 2017). To date, few studies have provided insights on the genetic basis of whitefish phenotypic traits (Vonlanthen *et al*. 2009; Gagnaire *et al*. 2013a; Gagnaire *et al*. 2013b; Feulner & Seehausen 2019; Jacobs *et al*. 2019). These studies focus mainly on trophic traits, whereas less attention has been paid on traits associated with timing of reproduction. Timing of reproduction can act as a reproductive barrier among diverging populations (Hendry & Day 2005), and it is well-known that whitefish morphs have differences in their spawning time and location (e.g. Svärdson (1979); Vonlanthen *et al*. (2009); Kahilainen *et al*. (2014); Bitz-Thorsen *et al*. (2020)).

Here, using a demographic modelling framework, we test two possible divergence scenarios of the Lake Tsahkal whitefish morphs: sympatric speciation and allopatric speciation followed by secondary contact. Furthermore, we investigate the genetic architecture of gonad weight variation, a proxy for spawning time which is a trait expected to be under ecological selection that could lead to reproductive isolation.

## Methods

### Study system and data collection

Lake Tsahkal is located in the treeline of northern Finnish Lapland (69°01’N, 20°50’E, 559 m a.s.l). The lake is oligotrophic (totP 5 µg/L, totN 140 µg/L), deep (max 35 m, mean 9 m), clearwater (compensation depth 7.5 m) and has equal habitat distribution (41% pelagic/profundal, 59% littoral; Hayden *et al*. (2014)). Introduced whitefish in the 1960’s had broad ecological opportunity, as they are more efficient in zooplanktivory and benthivory than native brown trouts (*Salmo trutta*) and burbots (*Lota lota*) (Siwertsson *et al*. 2010; Hayden *et al*. 2014). Reproductive isolation among whitefish morphs could arise via differences in resource use causing differences in spawning times and places (Kahilainen *et al*. 2014; Taylor & Friesen 2017; Thibert-Plante *et al*. 2020).

Littoral, pelagic and profundal zones were identified in the lake (Hayden *et al*. 2014) (Fig. 1a) and subsequently whitefish were collected from these habitats with gill net series in August 2011 (Kahilainen *et al*. 2011). Caught fish were transported to field laboratory, labelled and frozen (−20°C) for later measurements of fish size, gill raker count, resource use metrics and morphological traits (for details, see Table S1, Figure S2, S4; Method S1). All caught fish were assigned to three different morphs according to their gill raker appearance, head and body shape (Method S1; Kahilainen & Østbye (2006)).

**Figure 1.**
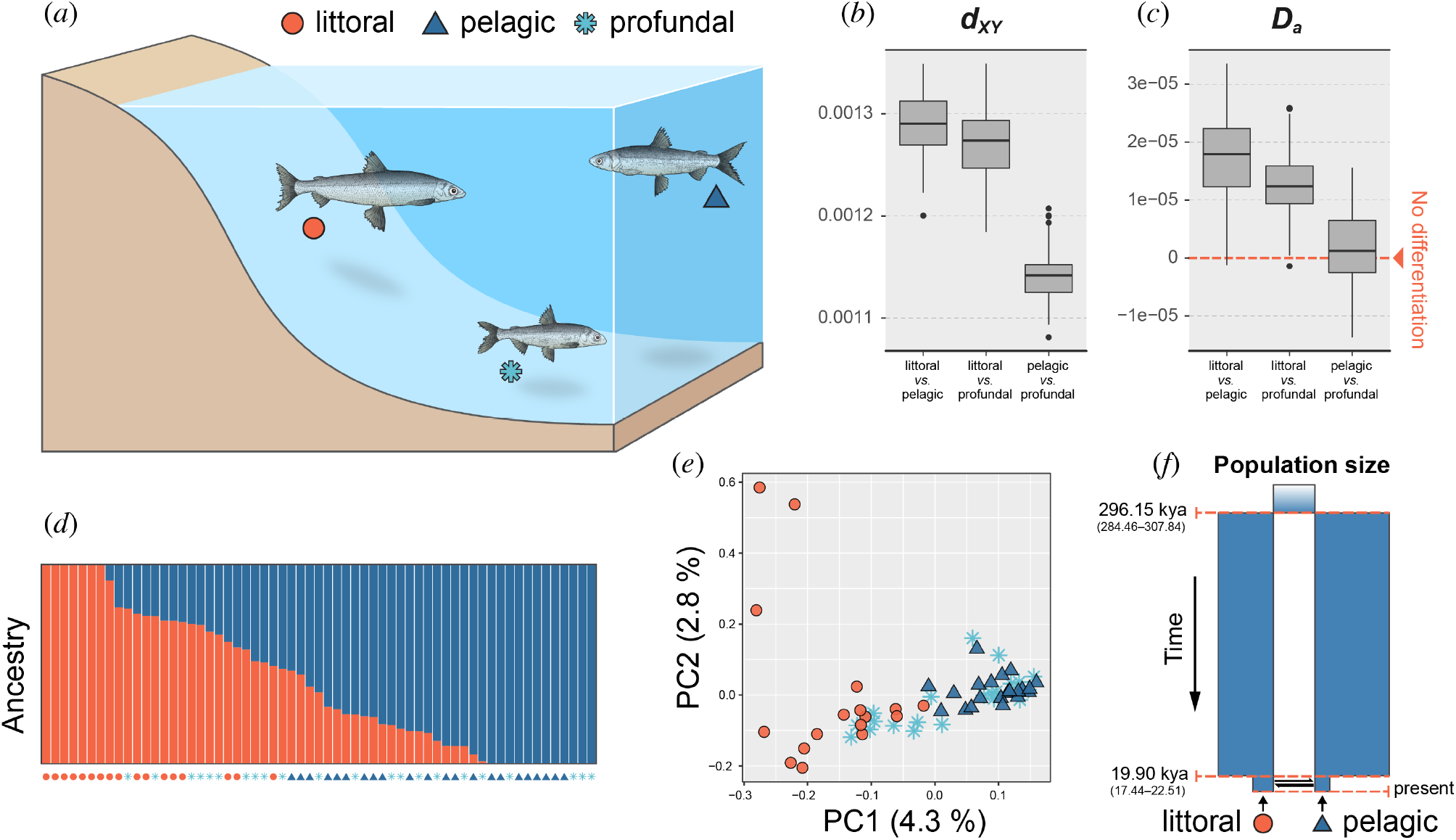
Population structure and demographic history of three morphs of the European whitefish (*Coregonus lavaretus*) in Lake Tsahkal. (a) Graphic presentation of the littoral, pelagic and profundal morphs inhabiting the lake. (b, c) Genome-wide divergence (absolute divergence, *d*_*XY*_ and net pairwise nucleotide divergence, *D*_*a*_) between morphs. Boxplots show 100 bootstraps by permuting sites within chromosome. (d) Individual ancestry reconstructed from *NGSadmix* at K = 2. (e) Principal component analysis (PCA) of genotype likelihoods of 15,615 SNPs. (f) Inferred demographic history of the littoral and pelagic morphs: secondary contact with changes in populations sizes. Estimated time parameters with 95% confidence intervals are shown. Kya, thousand years ago. Sample identifications of the (d,e) are presented in the electronic supplementary material, Figure S3.

A total of 61 individuals (17 littoral, 22 pelagic and 22 profundal fish) were sequenced using 2b-RAD sequencing approach (Wang *et al*. 2012) to obtain 36 bp single-end fragments with a mean coverage 30.2X (13.7–44.0; electronic supplementary material, Table S1) by Illumina HiSeq 4000 at BGI, Hong Kong. DNA extraction and 2b-RAD library preparation followed exactly the protocols detailed in Momigliano et al. (2018, 2021).

### Population genetic analyses

Raw reads were demultiplexed and PCR duplicates removed as per Momigliano *et al*. (2018). Reads were aligned to a chromosome-level assembly of the alpine European whitefish (*Coregonus* spp.) publicly available at https://www.ebi.ac.uk/ena/data/view/GCA_9021750, using *Bowtie2* (Langmead & Salzberg 2012). SAM files were converted to BAM files and indexed using *SAMtools* (Li & Durbin 2009).

Genotype likelihoods were estimated from BAM files using *ANGSD* (Korneliussen *et al*. 2014), retaining only biallelic loci with ≤ 25% missing data and bases with mapping quality and Phred scores >20. We performed a principal component analysis (PCA) using *PCAngsd* (Meisner & Albrechtsen 2018) retaining variants with minimum minor allele frequency of 0.02. Individual ancestries were inferred using *NGSadmix* (Skotte *et al*. 2012) based on genotype likelihoods, assuming 1-3 ancestral populations. Absolute divergence (*d*_*XY*_) (Nei 1987) and net nucleotide divergence (*D*_*a*_, Nei (1987)) were estimated based on non-admixed individuals identified by *NGSadmix* using scripts from Momigliano *et al*. (2021).

### Demographic modelling

We compared demographic models using the software package *moments* (Jouganous *et al*. 2017) based on diffusion approximations of the allele frequency spectrum (SFS). Ten demographic models were tested using non-admixed littoral and pelagic individuals, as the ancestral pelagic and profundal morphs were not genetically distinct (see Results) and we had more pelagic (n = 8) than profundal (n = 5) non-admixed individuals. We firstly defined five competing gene flow scenarios: strict isolation (SI), isolation with migration (IM), secondary contact (SC), ancient migration (AM) and a two epochs model (2EP) assuming heterogeneous migration rates through time (Roux *et al*. 2016; Momigliano *et al*. 2021). Since unaccounted changes in *N*_*e*_ can bias model choice and parameter estimation (Momigliano *et al*. 2021), we defined five additional models accounting for a change in *N*_*e*_ in both daughter populations at time T_2_, based on the above simple models: SI_NeC, AM_NeC, IM_NeC, SC_NeC and 2EP_NeC; NeC stands for “*N*_*e*_ Change”. The 10 tested models are visualized in electronic supplementary material, Figure S3. To account for possible effects of linkage in the 2b-RAD data, the best fitting model was chosen using Likelihood Ratio Test (LRT) as outlined in (Coffman *et al*. 2016).

All analyses were performed based on the unfolded two-dimensional SFS (2D-SFS) derived from the sites shared between pairwise morphs. The programs *ANGSD* and *Moments*, and custom R and python scripts adopted from Momigliano *et al*. (2021) and Fang *et al*. (2021) were used in the analyses. Detailed methods are given in the supplementary material, method S1.

### Admixture mapping

Admixture mapping of phenotypic traits was performed with two genome-wide association (GWA) approaches: GEMMA v.0.98, a genome-wide efficient mixed model association approach (Zhou & Stephens 2014), and LDna-EMMAX, a linkage disequilibrium (LD) clustering-based approach for association mapping (Kang *et al*. 2010; Kemppainen *et al*. 2015; Li *et al*. 2018; Fang *et al*. 2021). Univariate linear mixed models were conducted for each phenotypic trait in GEMMA, accounting for relatedness between individuals by supplying relatedness matrix as a covariate. In LDna-EMMAX, we used LD network analyses (LDna, Kemppainen *et al*. (2015)) to identify correlated clusters of single nucleotide polymorphisms (SNPs), followed by GWA by fitting a multilocus mixed model accounting for relatedness as in Li *et al*. (2018)) and in Fang et al. (2021). Significance tests of the two GWA approaches were performed based on Wald and permutation tests, respectively. Details and pipelines used perform GWAs are provided in the electronic supplementary material, method S1.

## Results

### (a) Population structure and demographic history

Littoral and pelagic morphs differed significantly in body size, gill raker count, morphology, diet and parasites, whereas profundal morph was intermediate (electronic supplementary material, Figure S4). Stable isotopes and mercury indicated clear differentiation of all three whitefish morphs in their respective habitats (electronic supplementary material, Figure S4). We obtained genotype likelihoods of 15,615 loci across 61 whitefish individuals. *NGSadmix* analyses revealed two ancestral components (K=2), and a clear genetic differentiation between littoral and pelagic/profundal morphs in analysis of non-admixed individuals (Fig. 1b-d). Overall divergence between non-admixed pelagic and profundal morphs was low: *d*_*XY*_=1.14e-3 (95%CI: 1.08e-3–1.21e-3; Fig. 1b) and *D*_*a*_= 1.81e-6 (95%CI: -1.37e-5–1.56e-5; Fig. 1c). Out of 61 individuals analysed, 41 were genetically admixed (i.e., having > 3% ancestry from a different cluster; Fig. 1d). PCA support these inferences, showing no differentiation of pelagic and profundal morphs, and admixture between littoral and pelagic morphs (Fig. 1e).

The demographic analyses identified secondary contact with changes in *N*_*e*_ (SC_NeC; Fig. 1f; electronic supplementary material, Figure S5, S6) as the best-fit model. Littoral and pelagic morphs diverged 296.15 thousand years ago (kya; 95% CI: 284.46–307.84 kya, i.e., 56.89–61.57 thousand generations), and experienced a secondary contact following the end of the last glaciation 19.9 kya (95% CI: 17.44–22.51 kya, 3.49–4.50 thousand generations) with decreases in *N*_*e*_ for both morphs after the Last Glacial Maximum (LGM; Fig. 1f). For a summary of scaled parameters, see the electronic supplementary material, Table S2.

### (b) Admixture mapping

Both GWA analyses revealed three SNPs (Chromosome [*Chr*] *22: 43,111,579, Chr 28: 9,855,214* and *Chr 36: 8,058,628*) associated with gonad weight variation in the 41 admixed individuals when treating age of individuals as a covariate in the models (Fig. 2a-c). The genotypes of the three SNPs explained 67.35% of the variation in gonad weight (*R*^2^=0.67; Fig. 2c). When using the body weight as a covariate, two of the three SNPs (*Chr 22: 43,111,579* and *Chr 36: 8,058,628*) were identified significantly associated with the gonad weight in the *GEMMA*, while not in LDna-EMMAX (electronic supplementary material, Fig. S8). However, these two SNPs are in strong LD (*r*^2^>0.9) and clustered in LDna (the first step in *LDna-EMMAX* analyses). No variant was significantly associated with any of the 22 other traits used to define the morphs (electronic supplementary material, Table S1).

**Figure 2.**
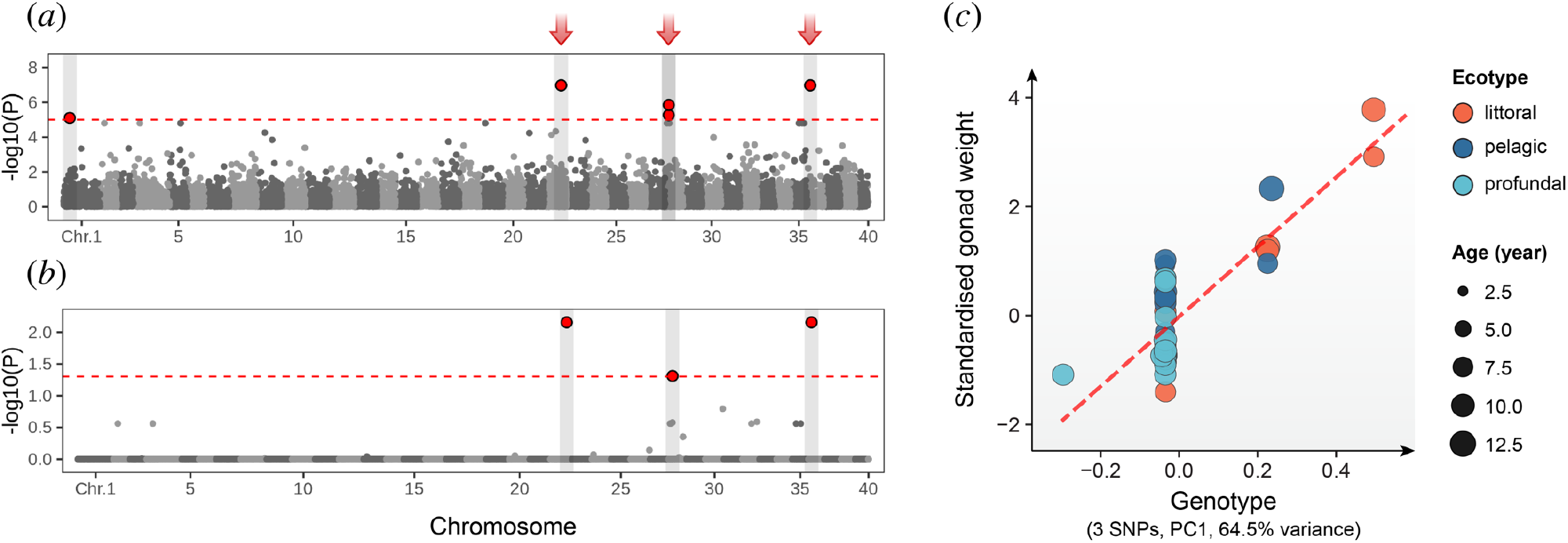
Admixture mapping of the gonad weight variation in European whitefish. Admixture mapping using *GEMMA* (a) and *LDna-EMMAX* (b) with age as covariate in models suggested three single-nucleotide polymorphisms (SNPs) significantly associated with the gonad weight variation. In (a,b,c), x-axis depicts genomic position and the y-axis the negative logarithm of the association *P*-value. Dashed red line marks the significance threshold adjusted for multiple testing. Chr = chromosome. (d) Correlation between the genotypes of gonad weight-associated SNPs and the standardised gonad weight, the residuals in a linear regression model relating the gonad weight to the body weight of individuals (y=6.4x-0.02; *R*^*2*^=0.67; *P*<5.1e-11).

## Discussion

Using demographic models, we refuted the hypothesis of rapid sympatric speciation of whitefish in Lake Tsahkal. Demographic analyses support an allopatric origin of whitefish morphs clearly predating the transplantation to the lake 50 years before sampling. This is not entirely surprising. While there are examples of rapid adaptive responses to novel environmental conditions on similar timescales (e.g. Bell *et al*. (2004); Lescak *et al*. (2015); Marques *et al*. (2019b)), in most cases these responses are subtle or have taken place over much longer time (Bolnick & Fitzpatrick 2007). For instance, the origins of most adaptive radiations in fishes can be traced back over hundreds of thousands years ago (e.g., Albertson *et al*. (1999); Danley & Kocher (2001); Hudson *et al*. (2007); Hudson *et al*. (2011); but see: Lescak *et al*. (2015); Jacobs *et al*. (2019).

Our results suggest that following a long phase of strict isolation, gene flow between divergent whitefish morphs was re-established during the last glacial retreat. Similar cases of secondary contact in post-glacial lakes have been reported from other northern temperate zone whitefish and they often led to hybridization (Bernatchez *et al*. 2010; Hudson *et al*. 2011; Rougeux *et al*. 2017; Rougeux *et al*. 2019b). The observed gradient of admixture suggests that the introduction of whitefish to the Lake Tsahkal created an artificial hybrid zone facilitating gene flow between littoral and pelagic morphs. The absence of genetic differentiation between the pelagic and profundal morphs was somewhat surprising. It is possible that these phenotypic differences are driven by plasticity but given our relatively low coverage of the whitefish genome it is also possible that we have missed important genetic differences between the two morphs.

An important finding from this study is the identification of three SNPs associated with gonad weight variation. Gonad weight is an interesting trait because of its association with spawning time and thereby also with reproductive isolation among whitefish morphs. It is known that whitefish morphs have different spawning times, habitats and size at maturity (Svärdson 1979; Vonlanthen *et al*. 2009; Bitz-Thorsen *et al*. 2020). The fact that the relative size of the gonads was largest in the littoral morph and smallest in the profundal morph makes perfectly sense as the former is an earlier breeder than the latter (Kahilainen *et al*. 2014).That differences in relative size of the gonads among the morphs were further mirrored in corresponding differences in frequency of three SNPs associated with gonad size strongly suggest that these SNPs are associated with breeding time differences among the morphs. Further weight on this inference is provided by the fact that in the lake whitefish (*C. clupeaformis*) gonadosomatic index maps to chromosome 28 similarly to one of the SNPs identified by us (Gagnaire *et al*. 2013a). Furthermore, two of the SNPs associated with gonad weight variation displayed strong long-range LD (*r*^2^>0.9) across chromosomes (*Chr. 22* and *36*), a pattern that could be indicative of strong selection (Lewontin & Kojima 1960; Nei & Li 1973). Ecological based selection for different spawning times may generate strong reproductive barriers and maintain reproductive isolation in syntopy as suggested by simulation models (Thibert-Plante *et al*. 2020) and empirical studies on European and Baltic flounders (Momigliano *et al*. 2017; Momigliano *et al*. 2018), the Atlantic cod (Fevolden & Pogson 1997; Hemmer-Hansen *et al*. 2013; Berg *et al*. 2016).

We found no SNP associated with gill raker number variation, a highly heritable trait playing a central role in adaptive radiations of coregonids (Bernatchez 2004; Rogers & Bernatchez 2007; Kahilainen *et al*. 2011). This is possibly a result of the small sample size of 41 admixed individuals and limited number of markers, which lowers the statistical power of GWA analyses (Hong & Park 2012). This could be particularly important if variation in gill raker number is a polygenic trait, which are notoriously difficult to map using GWA.

In conclusion, we provide strong evidence that the divergence of the littoral and pelagic morphs in Lake Tsahkal predates their introduction to the lake. While reproductive isolation among the two morphs is incomplete, their genetic differentiation suggests that spawning time differences have likely evolved in allopatry and the morphs are currently maintaining reproductive isolation in the absence of other clear barriers to gene flow. The identification of three SNPs associated with gonad weight variation, a proxy for spawning time, lends support to this hypothesis and provides a starting point to identify causal loci underlying reproductive isolation.

## Data accessibility

Information about samples used is provided in the supplementary material, Table S1. Raw sequence data has been uploaded to NCBI (PRJNA761129). The bioinformatic scripts used are deposited at Dryad (doi: xxx).

## Authors’ contributions

J.M. and P.M. conceive the study; B.F. conducted analyses with contributions from P.M.; B.F., J.M. and P.M. wrote the manuscript; K.K. provided samples, ecomorphological data and constructive comments on manuscript; Visualisation by BF; All authors approved the final version of the manuscript.

## Competing interests

Authors declare no competing interests.

## Funding

The research is supported by the Academy Finland (grants #129662, #134728, and #218343 to JM; #316294 to PM; #1140903 to KK), European Regional Development Fund (#A30205 to KK) and Finnish Cultural Foundation (# 00211290 to BF).

## Acknowledgments

We thank Tiina Holopainen and Petri Nieminen for their help in field work, Miinastiina Issakainen for help with laboratory work, Petri Kemppainen for advice and technical support in admixture mapping.

## Notes

### Competing Interest Statement

The authors have declared no competing interest.

