## Supplementary file for "Allopatric origin of sympatric whitefish morphs with insights on the genetic basis of their reproductive isolation"

### Supplementary Materials

**Figure S1. A picture illustrating the three different whitefish (*Coregonus lavaretus*) morphs.** Fish were classified according to their capturing habitat, i.e., littoral, pelagic or profundal, from top to bottom. Morphological measurements of 17 body traits were measured with accuracy of 0.001 mm. Also gill rakers were counted, gill arch length and longest gill raker length measured under preparation microscope from each sampled individual (Figure S2, Table S1, Method S1). Photo by Kimmo Kahilainen.

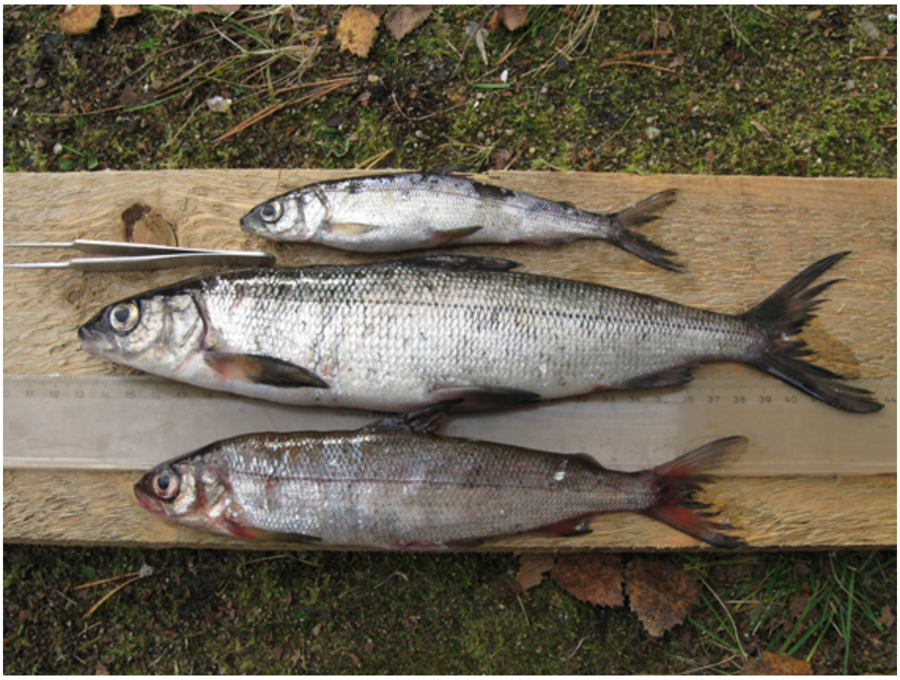

**Figure S2. The morphological traits measured in the study.** The linear morphometric measures are listed in the Table S1. Morphometric measures performed as per Vuorinen *et al.* (1993).

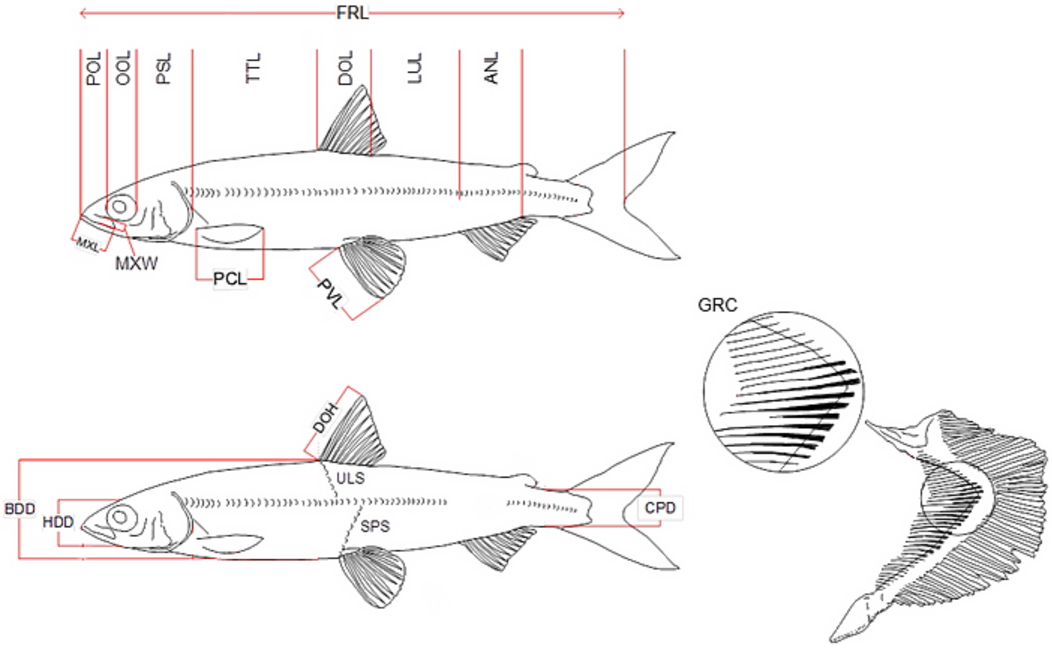

**Figure S3. Visualization of (a) individual ancestry and principal component analysis (PCA) of (b) marker data and (c) morphometric measures with individual identification.** The morphometric measures are corrected by length in PCA.

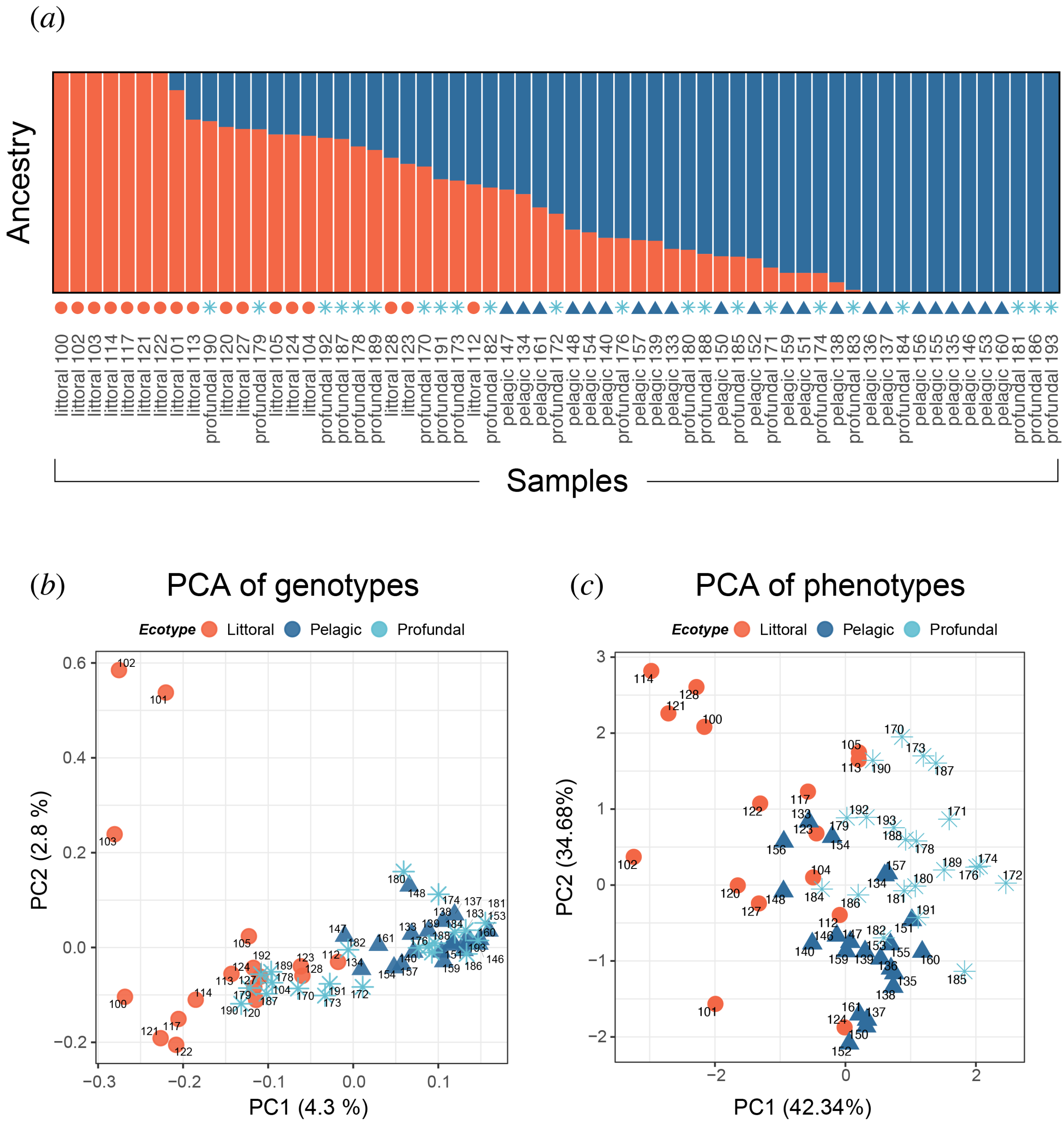

**Figure S4. Basic fish biological measurements (age, total length, weight, gonad weight), gill raker count, resource use metrics (pelagic parasites; *Diphyllobothrium* sp., diet, stable isotopes of carbon and nitrogen, total mercury) from whitefish morphs collected from littoral, pelagic and profundal habitats.** Gonad weight and mercury content was size or age corrected to mean total length (22.4 cm) or mean age (9.3 years) for all sampled individuals. Mercury content (µg/g) in in dry weight. C:N ratio refers to elemental ratio of carbon and nitrogen a proxy of lipid content in tissue. ANOVA and pairwise comparisons (Tukey's HSD test) was used to test statistical significance of measurements. For more details, see Method S1.

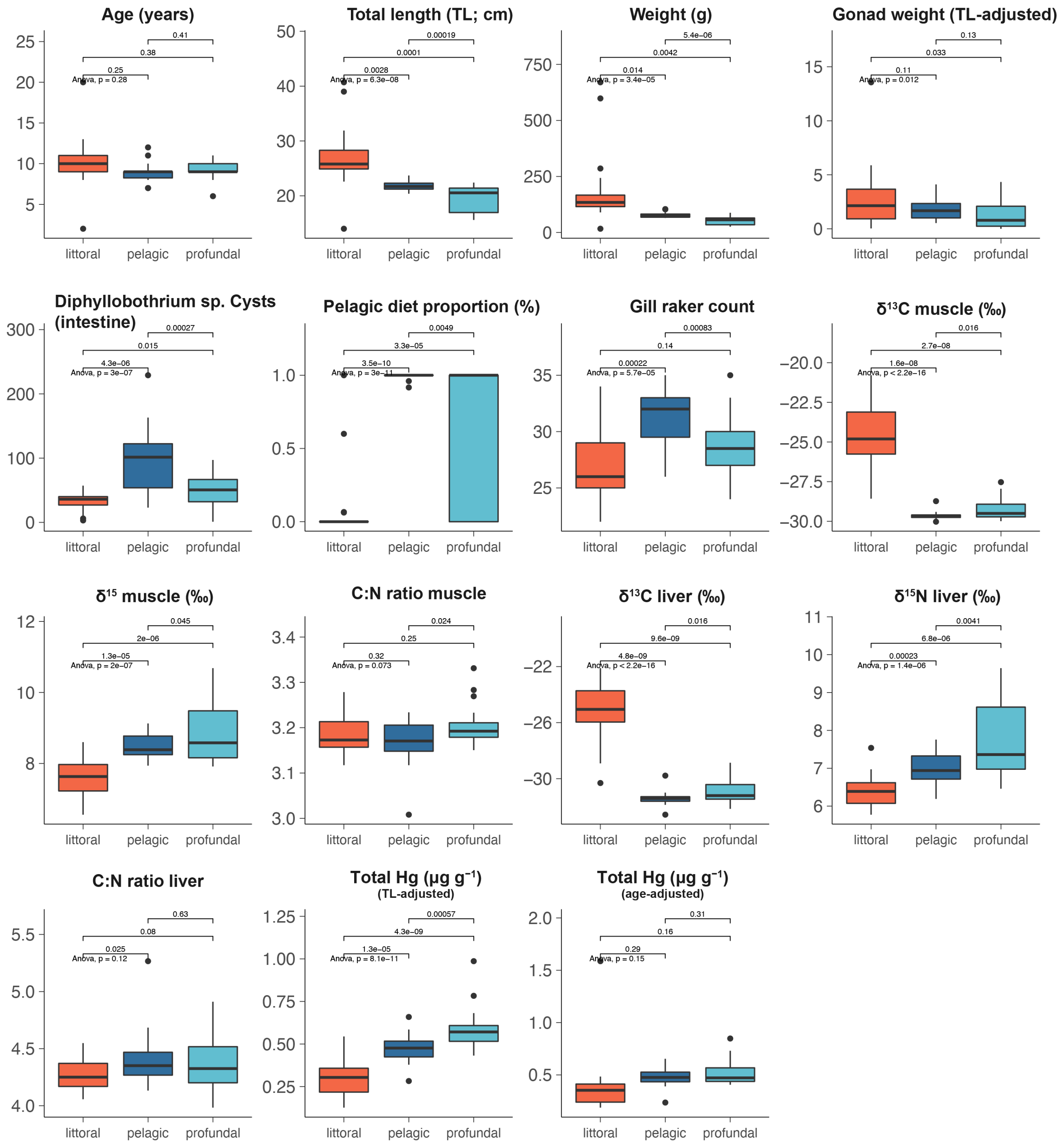

**Figure S5. Schematic representation of the 10 demographic models tested.**

The upper five plots present five simple models: strict isolation (SI), ancient migration (AM), isolation with migration (IM), secondary contact (SC) and two epochs (2EP) models. The bottom plots present five extended models incorporating changes in $N_e$ at $T_2$ based on the above five models; they are respectively SI_NeC, AM_NeC, IM_NeC, SC_NeC and 2EP_NeC.

For the first five models (SI, AM, IM, SC and 2EP), $N_{ref}$, ancestral population size; $N_{u1}$, current size of population 1; $N_{u2}$, current size of population 2; $T_1$, time of divergence; $T_2$, time at which migration stopped (in AM) or time of the secondary contact (in SC) or time at which migration rate changed (in 2EP); $m_{21}$, migration rate from population 1 to population 2; $m_{12}$, migration rate from population 2 to population 1.

For the five models in the bottom (SI_NeC, AM_NeC, IM_NeC, SC_NeC and 2EP_NeC), $T_1$, time of divergence; $T_2$, time of population size change and time at which migration stopped (in AM_NeC) or time of the secondary contact (in SC_NeC) or time at which migration rate changed (in 2EP_NeC); $N_{u1}$, size of population 1 before population size change ; $N_{u2}$, size of population 2 before population size change; $N_{u1b}$, current size of population 1; $N_{u2b}$, current size of population 2.

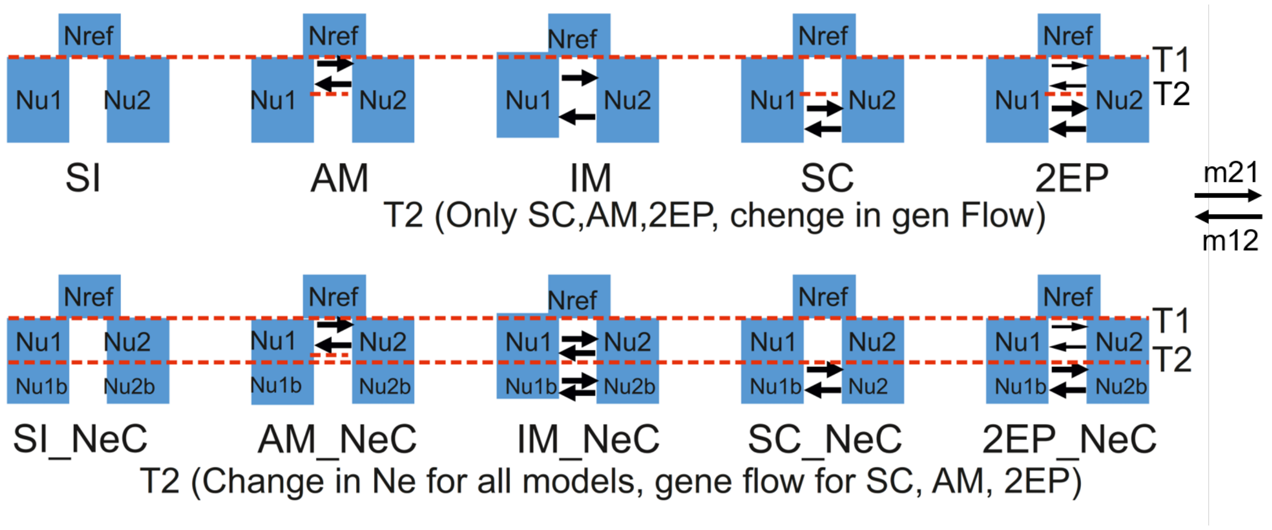

**Figure S6. The best-fit two-population model (SC-$N_e$C) and the two-dimensional allele frequency spectrum.** (a) Observed two-dimensional allele frequency spectrum (2D-SFS). (b) Modelled 2D-SFS based on SC_$N_e$C. (c-d) Modelled residuals.

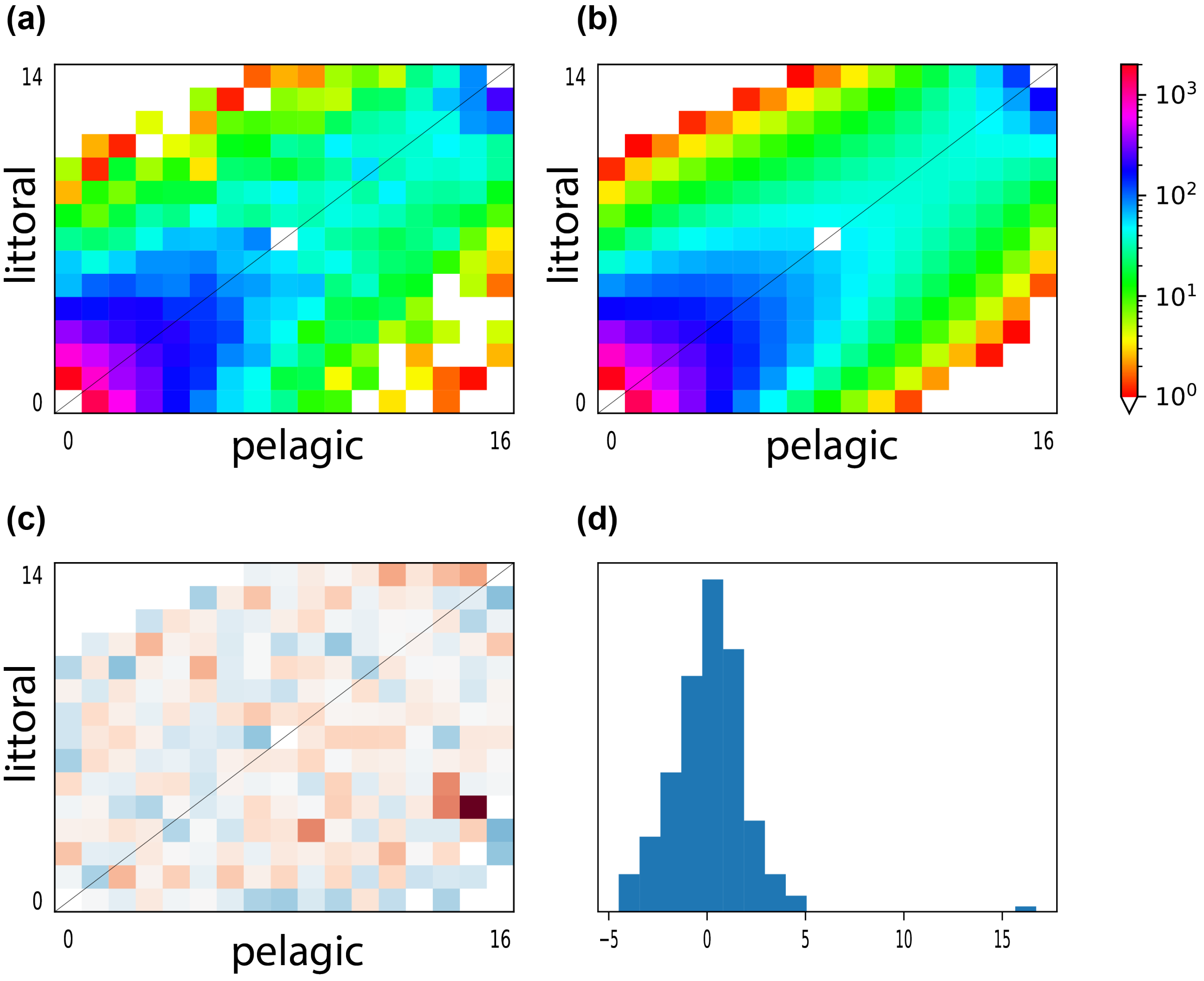

**Figure S7. Differences of gonad weight and the genotypes of the three associated SNPs among ecotypes.** The gonad weight and genotypes (PC 1 of the three SNPs) between ecotypes are compared for admixed individuals using analyses of variance (ANOVA) and subsequent Tukey's HSD pairwise tests. The standardised gonad weight is the residuals in a linear regression model relating the gonad weight to the body weight of individuals.

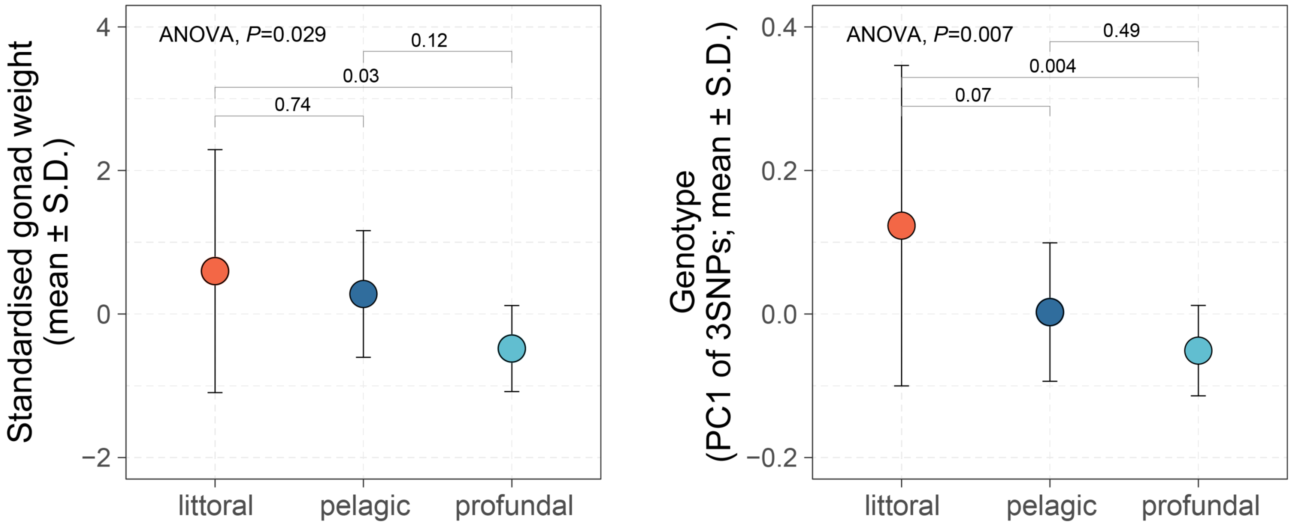

**Figure S8. Admixture mapping of the gonad weight variation.** Admixture mapping using *GEMMA* (a,b) and *EMMAX-LDna* (c,b) with age and body weight as covariate in models respectively. X-axis depicts genomic position and the y-axis the negative logarithm of the association *P*-value. Dashed red line marks the significance threshold adjusted for multiple testing. Chr = chromosome.

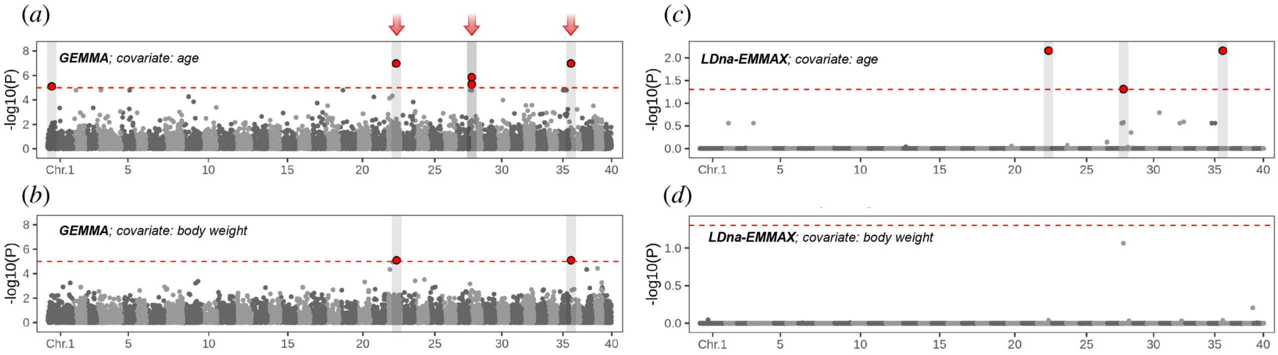

#### Table S1. Detailed sample information about sequencing depth, collection date and depth together with all linear morphometric measures.

*(This supplementary table will be prepared as a separate file in final sumission. )*

| Code | Capture habitat | Age | Sex | Sequencing depth | Admixed | Total length (cm) | Weight (g) | Gonad weight (g) | Gonad weight* | Diphyllobothrium sp. Cysts (intestine) | Pelagic diet proportion | d13C_muscle (‰) | d15N_muscle (‰) | C:N muscle |
|---|---|---|---|---|---|---|---|---|---|---|---|---|---|---|
| 11TSL100 | littoral | 10 | 2 | 16.1233 | No | 30.4 | 243.1 | 2.664 | 1.966452632 | 27 | 0.066 | -20.78874262 | 7.056566468 | 3.156935767 |
| 11TSL101 | littoral | 11 | 1 | 24.843 | Yes | 31.9 | 285.4 | 8.348 | 5.872386207 | 56 | 0 | -25.75970645 | 6.927341459 | 3.264076738 |
| 11TSL102 | littoral | 11 | 2 | 25.9093 | No | 39 | 598 | 7.046 | 4.05416 | 37 | 0 | -22.14412641 | 6.851272593 | 3.229578395 |
| 11TSL103 | littoral | 20 | 1 | 38.7188 | No | 40.7 | 670 | 24.603 | 13.5648973 | 40 | 0 | -22.0487904 | 6.557183249 | 3.278362938 |
| 11TSL104 | littoral | 9 | 2 | 27.3154 | Yes | 23.7 | 90.9 | 0.969 | 0.917483544 | 57 | 0 | -25.50039799 | 7.97054862 | 3.199460767 |
| 11TSL105 | littoral | 2 | 1 | 38.2011 | Yes | 14 | 16.7 | 0.022 | 0.035262857 | 36 | 1 | -28.56858553 | 7.75098798 | 3.174259344 |
| 11TSL112 | littoral | 9 | 1 | 44.0049 | Yes | 25.6 | 115.2 | 2.439 | 2.137935938 | 50 | 0.6 | -23.91443446 | 7.820461384 | 3.148303633 |
| 11TSL113 | littoral | 8 | 2 | 17.3859 | Yes | 25.1 | 118.2 | 0.808 | 0.722371315 | 47 | 0 | -24.8998058 | 7.542916156 | 3.155139469 |
| 11TSL114 | littoral | 13 | 2 | 38.8767 | No | 27.3 | 157.6 | 0.668 | 0.549081319 | 30 | 0 | -21.96779024 | 7.743777802 | 3.140639447 |
| 11TSL117 | littoral | 10 | 2 | 26.343 | No | 26.2 | 134.2 | 0.811 | 0.694612214 | 10 | 0 | -26.22205818 | 7.405573925 | 3.183747516 |
| 11TSL120 | littoral | 12 | 1 | 26.4846 | Yes | 25.6 | 124.3 | 3.555 | 3.116179688 | 6 | 0 | -27.15888907 | 8.598653967 | 3.160058314 |
| 11TSL121 | littoral | 10 | 1 | 20.4451 | No | 28.3 | 166.5 | 4.318 | 3.423884099 | 3 | 0 | -25.17693407 | 8.088919807 | 3.117398372 |
| 11TSL122 | littoral | 9 | 1 | 28.3169 | No | 22.9 | 102.3 | 3.739 | 3.66389345 | 27 | 0 | -26.40192367 | 8.259756277 | 3.158195841 |
| 11TSL123 | littoral | 9 | 2 | 40.4246 | Yes | 22.6 | 89.9 | 1.886 | 1.872647788 | 38 | 0 | -24.59336374 | 7.629921418 | 3.256678071 |
| 11TSL124 | littoral | 10 | 1 | 19.1348 | Yes | 26.2 | 140 | 3.71 | 3.177572519 | 35 | 0.0625 | -24.80494693 | 7.610598447 | 3.213279593 |
| 11TSL127 | littoral | 8 | 1 | 25.2233 | Yes | 24.9 | 129.9 | 5.289 | 4.766472289 | 37 | 0 | -23.64390871 | 8.425441046 | 3.172866156 |
| 11TSL128 | littoral | 9 | 2 | 21.9212 | Yes | 25.8 | 137.9 | 1.992 | 1.732576744 | 13 | 0 | -23.10711438 | 7.225127864 | 3.164850851 |
| 11TSL133 | pelagic | 8 | 1 | 18.9607 | Yes | 23.4 | 104.3 | 2.543 | 2.438671795 | 41 | 0.96 | -28.72808215 | 8.359545182 | 3.164547737 |
| 11TSL134 | pelagic | 7 | 1 | 41.8016 | Yes | 22.3 | 80.1 | 1.375 | 1.383632287 | 117 | 1 | -29.50449579 | 8.500356478 | 3.155503949 |
| 11TSL135 | pelagic | 9 | 2 | 14.7916 | No | 22.3 | 82.1 | 1.1 | 1.10690583 | 106 | 1 | -29.65251151 | 8.150756584 | 3.205954504 |
| 11TSL136 | pelagic | 9 | 2 | 22.6934 | No | 21.3 | 67 | 1.023 | 1.077752113 | 102 | 1 | -29.65967324 | 8.411668579 | 3.117313038 |
| 11TSL137 | pelagic | 8 | 2 | 32.3172 | No | 21.6 | 73.4 | 0.541 | 0.562038889 | 104 | 1 | -29.74416125 | 8.333682884 | 3.169182677 |
| 11TSL138 | pelagic | 7 | 1 | 37.6142 | Yes | 21.7 | 72.5 | 2.498 | 2.583185253 | 77 | 1 | -29.98560492 | 8.244301173 | 3.164547737 |
| 11TSL139 | pelagic | 9 | 1 | 39.5013 | Yes | 22 | 81.2 | 1.92 | 1.9584 | 99 | 1 | -29.75848471 | 8.02481037 | 3.171976652 |
| 11TSL140 | pelagic | 9 | 2 | 34.2586 | Yes | 23.7 | 103.1 | 1.126 | 1.066136709 | 96 | 1 | -29.75666035 | 8.966292996 | 3.162233168 |
| 11TSL146 | pelagic | 9 | 1 | 25.8569 | No | 21 | 74.7 | 1.91 | 2.040971429 | 34 | 1 | -29.51795936 | 7.939591535 | 3.145652916 |
| 11TSL147 | pelagic | 9 | 1 | 24.1994 | Yes | 21.1 | 71.2 | 2.563 | 2.72576872 | 84 | 1 | -29.69748709 | 8.530780454 | 3.130737307 |
| 11TSL148 | pelagic | 10 | 1 | 35.1959 | Yes | 21.4 | 68.2 | 1.939 | 2.033231776 | 132 | 1 | -29.49531227 | 8.471423917 | 3.008063914 |
| 11TSL150 | pelagic | 11 | 2 | 41.1945 | Yes | 22 | 81.6 | 0.965 | 0.9843 | 229 | 1 | -29.75208176 | 8.955295779 | 3.136016784 |
| 11TSL151 | pelagic | 10 | 2 | 17.6939 | Yes | 22.7 | 82.2 | 0.991 | 0.979649339 | 102 | 0.916666667 | -29.62159235 | 8.835888893 | 3.131088698 |
| 11TSL152 | pelagic | 8 | 1 | 38.3217 | Yes | 21.2 | 68.5 | 2.466 | 2.610237736 | 137 | 1 | -29.97783942 | 9.126826063 | 3.219728442 |
| 11TSL153 | pelagic | 9 | 2 | 27.534 | No | 21.6 | 72.9 | 1.215 | 1.26225 | 124 | 1 | -29.69003276 | 8.116672321 | 3.198719505 |
| 11TSL154 | pelagic | 9 | 1 | 29.7387 | Yes | 22.5 | 83.2 | 2.058 | 2.052512 | 163 | 1 | -29.60435238 | 8.918037464 | 3.189969489 |
| 11TSL155 | pelagic | 9 | 2 | 23.4794 | No | 21.9 | 82.3 | 0.871 | 0.892476712 | 42 | 1 | -29.97923647 | 8.908722886 | 3.230791763 |
| 11TSL156 | pelagic | 9 | 1 | 24.9335 | No | 20.9 | 66.2 | 2.665 | 2.861368421 | 101 | 1 | -29.68044917 | 8.288896418 | 3.215717633 |
| 11TSL157 | pelagic | 9 | 1 | 30.05 | Yes | 20.8 | 66.9 | 1.817 | 1.960263462 | 35 | 1 | -29.40829831 | 7.939296524 | 3.204800901 |
| 11TSL159 | pelagic | 9 | 2 | 34.0516 | Yes | 21.7 | 65.2 | 0.875 | 0.90483871 | 46 | 1 | -29.65390128 | 8.581221213 | 3.233763667 |
| 11TSL160 | pelagic | 8 | 2 | 29.183 | No | 20.4 | 66.2 | 0.484 | 0.5324 | 23 | 1 | -29.59683069 | 8.351722019 | 3.196278847 |
| 11TSL161 | pelagic | 12 | 1 | 25.5018 | Yes | 22.6 | 89.1 | 4.13 | 4.100761062 | 163 | 1 | -30.022964 | 8.256494793 | 3.207959475 |
| 11TSL170 | profundal | 9 | 2 | 31.9665 | Yes | 16 | 26.7 | 0.027 | 0.0378675 | 26 | 0 | -27.94618299 | 9.763997852 | 3.186098775 |
| 11TSL171 | profundal | 6 | 1 | 35.0188 | Yes | 15.6 | 25.9 | 0.125 | 0.179807692 | 60 | 0 | -28.85251238 | 9.314314042 | 3.209903862 |
| 11TSL172 | profundal | 9 | 1 | 34.31 | Yes | 16.6 | 29.2 | 0.112 | 0.15140241 | 24 | 0 | -28.44359684 | 9.515175687 | 3.182561868 |
| 11TSL173 | profundal | 9 | 2 | 31.4488 | Yes | 18 | 40 | 0.027 | 0.03366 | 52 | 0 | -28.45282048 | 9.95832017 | 3.150517122 |
| 11TSL174 | profundal | 9 | 1 | 39.5611 | Yes | 15.7 | 26.2 | 0.114 | 0.162940127 | 78 | 0.966666667 | -29.41792077 | 9.768854542 | 3.195715976 |
| 11TSL176 | profundal | 10 | 1 | 30.2266 | Yes | 16.5 | 32.8 | 0.353 | 0.48008 | 31 | 0 | -29.43534687 | 9.379914895 | 3.169464883 |
| 11TSL178 | profundal | 9 | 2 | 40.0494 | Yes | 19.6 | 54.6 | 0.548 | 0.627404082 | 35 | 1 | -29.72019724 | 8.320413008 | 3.162250598 |
| 11TSL179 | profundal | 9 | 1 | 34.505 | Yes | 18.9 | 53.2 | 1.98 | 2.350857143 | 1 | 0 | -27.52838057 | 10.68308724 | 3.269373438 |
| 11TSL180 | profundal | 9 | 2 | 27.0833 | Yes | 21.2 | 65.4 | 0.38 | 0.402226415 | 35 | 1 | -29.43536279 | 8.782186648 | 3.180747965 |
| 11TSL181 | profundal | 9 | 2 | 28.4565 | No | 21.4 | 67.4 | 0.665 | 0.697317757 | 58 | 1 | -29.56609239 | 8.502645496 | 3.17597599 |
| 11TSL182 | profundal | 9 | 1 | 35.5422 | Yes | 21.3 | 63 | 2.115 | 2.228197183 | 16 | 1 | -29.71656709 | 8.113012035 | 3.207681873 |
| 11TSL183 | profundal | 11 | 1 | 29.4422 | No | 21.4 | 65.1 | 4.13 | 4.33071028 | 38 | 1 | -29.98684217 | 8.153739415 | 3.211227922 |
| 11TSL184 | profundal | 11 | 1 | 37.8031 | No | 21.8 | 87.7 | 3.874 | 3.98773211 | 97 | 0.928571429 | -29.12346723 | 7.994298961 | 3.283170958 |
| 11TSL185 | profundal | 9 | 1 | 42.5741 | Yes | 21.4 | 62.4 | 2.047 | 2.146480374 | 49 | 1 | -29.8082142 | 7.91700708 | 3.199935548 |
| 11TSL186 | profundal | 8 | 1 | 37.0259 | No | 19.6 | 52.2 | 1.936 | 2.216522449 | 42 | 1 | -29.86240113 | 9.038640755 | 3.180322354 |
| 11TSL187 | profundal | 9 | 2 | 36.3173 | Yes | 15.7 | 25.8 | 0.002 | 0.002858599 | 63 | 0 | -28.38724954 | 9.970114989 | 3.232883529 |
| 11TSL188 | profundal | 9 | 2 | 31.2565 | Yes | 21.6 | 56.8 | 0.837 | 0.86955 | 22 | 1 | -29.83878569 | 8.659709499 | 3.1783768 |
| 11TSL189 | profundal | 9 | 1 | 13.7311 | Yes | 20.6 | 57.5 | 1.326 | 1.444438835 | 72 | 1 | -29.57473445 | 8.180000514 | 3.164312779 |
| 11TSL190 | profundal | 10 | 1 | 17.6318 | Yes | 20.5 | 55.7 | 0.594 | 0.650212683 | 68 | 1 | -29.39907178 | 8.300795026 | 3.193556991 |
| 11TSL191 | profundal | 10 | 2 | 41.7556 | Yes | 21.4 | 59.6 | 0.94 | 0.985682243 | 83 | 1 | -29.6490583 | 8.06128744 | 3.191127208 |
| 11TSL192 | profundal | 10 | 2 | 21.2357 | Yes | 22.4 | 83.5 | 1.07 | 1.071910714 | 72 | 1 | -29.6727199 | 8.071989644 | 3.229301584 |
| 11TSL193 | profundal | 10 | 1 | 37.2179 | No | 21.2 | 68.2 | 1.784 | 1.88834717 | 57 | 1 | -29.81900508 | 8.422977164 | 3.331222212 |

| Code | δ13C liver (‰) | δ15N liver (‰) | C:N liver | Mean THg (mg/kg) | THg (mg/k)* | THg (mg/k)** | Gill raker count (GRC) | Gill Arch Length* | Caudal peduncle depth (CPD)* | Anal fin length (ANL)* | Lumbar length (LUL)* | Dorsal length (DOL)* | Trunk length (TTL)* |
|---|---|---|---|---|---|---|---|---|---|---|---|---|---|
| 11TSL100 | -22.11904181 | 5.855200046 | 4.249865529 | 0.25643595 | 0.189290221 | 0.239511177 | 23 | 23.46688056 | 23.05604777 | 19.07531898 | 42.07183552 | 20.24731063 | 51.39710487 |
| 11TSL101 | -24.87372943 | 6.234399534 | 4.547893363 | 0.2550387 | 0.179406534 | 0.216551042 | 31 | 24.15131123 | 22.82986891 | 21.66281256 | 39.42132941 | 18.34015451 | 46.52379075 |
| 11TSL102 | -22.62437416 | 6.274025768 | 4.162439712 | 0.22011905 | 0.126653115 | 0.186901084 | 26 | 23.60304079 | 27.73439091 | 20.74771712 | 38.23797205 | 18.86657268 | 47.54206982 |
| 11TSL103 | -22.90433094 | 5.87466161 | 4.38083549 | 0.5511271 | 0.303864671 | 0.257376356 | 33 | 27.45682526 | 23.75351279 | 20.16900665 | 35.68079065 | 18.12450502 | 49.81081647 |
| 11TSL104 | -25.10364719 | 6.972142129 | 4.370998651 | 0.45784655 | 0.433505341 | 0.475142975 | 29 | 25.41773097 | 29.23648993 | 19.22623447 | 35.88875034 | 18.33639095 | 49.75696201 |
| 11TSL105 | -30.30767331 | 6.61750071 | 4.244976117 | 0.33978115 | 0.544620643 | 1.586777971 | 25 | 25.82458448 | 25.1010887 | 21.16193205 | 38.42368782 | 18.10268205 | 48.61839388 |
| 11TSL112 | -25.99543505 | 6.505920356 | 4.180060945 | 0.46593485 | 0.408421017 | 0.483536833 | 30 | 23.49734936 | 25.17884722 | 20.19614749 | 35.79339332 | 19.4167137 | 48.76741294 |
| 11TSL113 | -24.71733865 | 6.387755725 | 4.434105867 | 0.30776295 | 0.275147434 | 0.359313244 | 25 | 21.6247741 | 26.19351211 | 16.08447374 | 39.59942434 | 21.28275607 | 41.01334769 |
| 11TSL114 | -22.12411569 | 6.486417313 | 4.168408225 | 0.3988813 | 0.327871662 | 0.286580872 | 22 | 26.32615944 | 23.65699427 | 21.18253506 | 37.89335067 | 20.84347173 | 44.2077589 |
| 11TSL117 | -28.60681941 | 5.773949794 | 4.427099154 | 0.2947178 | 0.252422421 | 0.275266425 | 26 | 23.51135116 | 25.74781515 | 16.84505838 | 41.56737504 | 17.56430259 | 45.37816567 |
| 11TSL120 | -28.90588181 | 7.537496449 | 4.14305934 | 0.2484188 | 0.217754604 | 0.193352633 | 28 | 25.76005708 | 24.46489623 | 17.36566282 | 39.90149771 | 19.33658176 | 45.85611273 |
| 11TSL121 | -25.61984109 | 6.330536019 | 4.09386997 | 0.2265177 | 0.179613328 | 0.211567532 | 22 | 23.95899853 | 22.7711259 | 22.41969468 | 36.70438638 | 19.13790198 | 46.61880453 |
| 11TSL122 | -25.95643223 | 6.764146538 | 4.18779321 | 0.36529395 | 0.357956168 | 0.379093944 | 25 | 24.11417767 | 22.50397683 | 20.35561728 | 36.99835573 | 19.23490357 | 51.7309543 |
| 11TSL123 | -25.05407265 | 6.609391768 | 4.280373798 | 0.43061945 | 0.427570817 | 0.446887296 | 27 | 24.21420358 | 22.3899425 | 23.61267361 | 36.16107321 | 19.84398941 | 47.88981844 |
| 11TSL124 | -25.53286111 | 6.073203534 | 4.281648225 | 0.3776404 | 0.323444678 | 0.352716134 | 34 | 24.02246749 | 20.45484526 | 20.97184425 | 39.76851046 | 19.19431094 | 47.73619106 |
| 11TSL127 | -23.73109354 | 6.751651179 | 4.25581369 | 0.33441825 | 0.301379339 | 0.390433307 | 28 | 24.28496627 | 22.4021824 | 17.25991433 | 42.58785127 | 18.56749061 | 48.15062165 |
| 11TSL128 | -24.80725743 | 5.920008887 | 4.056593994 | 0.39721295 | 0.345482891 | 0.412218773 | 22 | 24.71164421 | 25.44122448 | 21.26690927 | 31.13997308 | 21.88108454 | 44.50387491 |
| 11TSL133 | -29.78090524 | 6.688392837 | 4.200801797 | 0.395031 | 0.3788246 | 0.461198693 | 26 | 23.66252966 | 23.50197066 | 18.10106261 | 40.19497914 | 20.51605584 | 44.5295559 |
| 11TSL134 | -31.03137886 | 7.176150695 | 4.194507456 | 0.46419835 | 0.4671126 | 0.619373227 | 29 | 23.05594781 | 24.4625746 | 18.05732988 | 31.78580402 | 19.53192667 | 40.78304853 |
| 11TSL135 | -31.50260071 | 7.004836135 | 4.50125366 | 0.5531648 | 0.556637583 | 0.574062137 | 33 | 24.44128373 | 24.65621502 | 18.68175691 | 37.30099999 | 20.72631089 | 51.41921359 |
| 11TSL136 | -31.32799 | 7.260235482 | 4.34027899 | 0.4891916 | 0.515373686 | 0.507672172 | 33 | 24.78456609 | 29.76452517 | 19.36944975 | 36.11793666 | 20.44390798 | 45.03595174 |
| 11TSL137 | -31.4244661 | 7.731501265 | 4.399644541 | 0.46923545 | 0.487483495 | 0.547832388 | 35 | 26.73115609 | 25.38362013 | 21.14936413 | 32.32859548 | 21.00766016 | 46.99730748 |
| 11TSL138 | -31.28748392 | 7.346304616 | 4.132754221 | 0.48970595 | 0.5064056 | 0.653407653 | 33 | 23.69151456 | 23.28774455 | 19.21751429 | 39.79605205 | 20.59043853 | 44.76676632 |
| 11TSL139 | -31.36056695 | 6.70998493 | 4.348869213 | 0.50763975 | 0.517792545 | 0.526817252 | 32 | 24.12043496 | 26.70598778 | 20.72461234 | 37.84493189 | 21.0568012 | 47.93320347 |
| 11TSL140 | -31.55892433 | 7.214074194 | 4.419258274 | 0.4202779 | 0.397934012 | 0.436155065 | 31 | 26.51329122 | 27.29774624 | 21.6826361 | 34.5767625 | 20.22314948 | 47.3862282 |
| 11TSL146 | -31.20961947 | 6.390883955 | 4.244073509 | 0.44305595 | 0.473436929 | 0.459793619 | 31 | 25.49793568 | 23.77968638 | 23.57465852 | 34.90930512 | 20.35414677 | 49.0336212 |
| 11TSL147 | -31.28560043 | 6.719023408 | 4.187549254 | 0.50516225 | 0.537243644 | 0.524246157 | 31 | 24.56711891 | 24.46037446 | 21.73855141 | 35.79471387 | 19.93245744 | 45.48314847 |
| 11TSL148 | -31.25553416 | 6.531577006 | 4.35684501 | 0.55775665 | 0.584862581 | 0.520944711 | 29 | 26.53110503 | 24.7005519 | 19.90026739 | 36.76798548 | 19.60854065 | 46.9833274 |
| 11TSL150 | -31.86074732 | 7.032510901 | 4.506853156 | 0.27727445 | 0.282819939 | 0.235431215 | 35 | 26.42238518 | 22.50536054 | 22.47019073 | 38.73773238 | 23.99225865 | 42.1766963 |
| 11TSL151 | -31.54373798 | 7.368652968 | 4.46318556 | 0.4574627 | 0.452223039 | 0.427270162 | 31 | 23.00665221 | 23.6466734 | 20.06294031 | 35.79719703 | 20.91231214 | 50.92304514 |
| 11TSL152 | -31.71598765 | 7.743237052 | 4.352914609 | 0.37089975 | 0.392593886 | 0.433025458 | 35 | 26.22495417 | 23.94747981 | 17.25874224 | 40.03148606 | 20.49490565 | 44.37317106 |
| 11TSL153 | -31.6388485 | 6.748415526 | 4.469573583 | 0.4086672 | 0.424559813 | 0.424105739 | 32 | 25.20656544 | 26.92502995 | 17.72871836 | 37.26990486 | 21.05471208 | 47.86742189 |
| 11TSL154 | -31.40237126 | 6.722270645 | 4.343525797 | 0.4260225 | 0.42488644 | 0.442116683 | 27 | 23.66937944 | 25.02936405 | 16.23149867 | 38.38846255 | 20.4137626 | 47.70156305 |
| 11TSL155 | -31.81392642 | 7.583801609 | 4.683802797 | 0.37605175 | 0.385324259 | 0.390258149 | 32 | 24.41385504 | 23.8946714 | 25.23791477 | 32.19789369 | 22.33067226 | 50.16332353 |
| 11TSL156 | -30.99847005 | 6.64843677 | 4.172904475 | 0.47220225 | 0.5069961 | 0.490041002 | 27 | 25.18241094 | 25.47946856 | 18.56000246 | 39.21461122 | 19.47085391 | 47.5806055 |
| 11TSL157 | -31.28337083 | 6.190153165 | 4.342600863 | 0.6108878 | 0.659053953 | 0.633965784 | 29 | 22.87106755 | 19.96702185 | 20.59899089 | 34.55000953 | 20.08264795 | 45.1743921 |
| 11TSL159 | -31.5862266 | 7.756369585 | 4.622464219 | 0.42590525 | 0.440429208 | 0.441995004 | 32 | 26.12141349 | 23.08530342 | 23.26497397 | 31.70269537 | 22.91002942 | 44.74356065 |
| 11TSL160 | -32.56487857 | 6.875796225 | 5.266282077 | 0.43615255 | 0.479767805 | 0.509208102 | 33 | 24.10028858 | 29.40612979 | 16.95493986 | 37.9292754 | 22.48973963 | 46.71379615 |
| 11TSL161 | -31.61462651 | 6.822874426 | 4.343747114 | 0.52691565 | 0.523185274 | 0.410116014 | 33 | 23.87491763 | 26.23425193 | 19.58562045 | 37.7038468 | 21.23387483 | 43.12805816 |
| 11TSL170 | -29.99218098 | 9.642350527 | 4.566284459 | 0.7035149 | 0.986679647 | 0.73009213 | 24 | 23.62125428 | 24.67564284 | 23.80824375 | 32.26763013 | 20.70945163 | 48.20353213 |
| 11TSL171 | -30.36736921 | 8.627725265 | 4.152924748 | 0.54448535 | 0.783221234 | 0.847582195 | 27 | 23.40635308 | 24.84861079 | 16.60552825 | 39.81849695 | 19.64265054 | 45.10201302 |
| 11TSL172 | -29.74108846 | 9.094875538 | 4.102049512 | 0.39092885 | 0.528460445 | 0.405697273 | 30 | 22.04753435 | 25.4109476 | 21.82398869 | 32.22971736 | 21.1000292 | 44.7215475 |
| 11TSL173 | -30.56781389 | 9.220132389 | 4.512873349 | 0.54650025 | 0.681303645 | 0.567145815 | 25 | 22.44767166 | 28.31543378 | 17.83816373 | 33.70571773 | 20.38571649 | 44.04853613 |
| 11TSL174 | -30.81168518 | 8.573558999 | 3.983104804 | 0.41998485 | 0.600284079 | 0.435850944 | 29 | 23.04890666 | 26.48632572 | 20.66830479 | 33.62827018 | 19.53511896 | 44.23518676 |
| 11TSL176 | -31.29710483 | 8.50734527 | 4.319420013 | 0.45701805 | 0.621544548 | 0.426854859 | 29 | 22.63001459 | 25.33283825 | 19.90929627 | 34.1135928 | 23.22653649 | 47.19498097 |
| 11TSL178 | -31.53769992 | 7.297513281 | 4.532219979 | 0.43497695 | 0.498004222 | 0.451409413 | 28 | 22.99630297 | 25.77999562 | 21.69805611 | 34.89890999 | 22.21825371 | 44.11151451 |
| 11TSL179 | -28.86324265 | 9.441672198 | 4.48488164 | 0.41335405 | 0.49077592 | 0.428969647 | 26 | 23.9213133 | 27.35366595 | 17.92411882 | 40.08164814 | 24.68423928 | 46.13347154 |
| 11TSL180 | -31.48521261 | 7.196829203 | 4.583659796 | 0.55075955 | 0.582973788 | 0.571566022 | 30 | 23.31096889 | 23.90966138 | 23.64410786 | 33.73950703 | 20.58808661 | 48.67411831 |
| 11TSL181 | -31.5059852 | 7.244595869 | 4.518341532 | 0.56707255 | 0.594631216 | 0.588495291 | 30 | 23.46474942 | 24.23179055 | 22.14312664 | 37.18812944 | 19.18235398 | 46.29253738 |
| 11TSL182 | -30.91191329 | 6.459783091 | 4.074983235 | 0.5637175 | 0.593888296 | 0.585013494 | 31 | 23.74629836 | 25.57650606 | 20.87205921 | 34.2008358 | 18.05667595 | 45.15854711 |
| 11TSL183 | -31.41775644 | 7.669226983 | 4.100103768 | 0.51490225 | 0.539925537 | 0.43719882 | 35 | 24.38293527 | 26.11948662 | 20.22620966 | 34.09778302 | 20.41455023 | 45.55988092 |
| 11TSL184 | -30.08934916 | 6.747603735 | 4.330252214 | 0.5525558 | 0.568777622 | 0.469170107 | 28 | 24.06978256 | 22.69355942 | 22.23747876 | 37.52111686 | 21.42834452 | 48.04911218 |
| 11TSL185 | -31.29102691 | 6.475968075 | 4.261359954 | 0.5465999 | 0.573163633 | 0.56724923 | 33 | 21.42433643 | 23.63383112 | 20.80427832 | 35.66766253 | 22.24330226 | 45.53408315 |
| 11TSL186 | -31.08369929 | 7.583303721 | 4.1807772 | 0.46667045 | 0.534290046 | 0.54483775 | 29 | 24.62339988 | 25.90451479 | 23.47190348 | 35.15882062 | 18.73347714 | 48.24849579 |
| 11TSL187 | -29.11384078 | 9.317546511 | 4.277164356 | 0.4277554 | 0.611390521 | 0.443915048 | 25 | 23.04890666 | 19.95003146 | 22.869568 | 35.90423804 | 21.19418202 | 45.61463834 |
| 11TSL188 | -31.36236221 | 7.426563291 | 4.473608552 | 0.4163147 | 0.432504716 | 0.432042144 | 28 | 22.40064141 | 22.77632719 | 21.02516307 | 30.65163547 | 20.74054767 | 42.9823297 |
| 11TSL189 | -31.11445299 | 6.569705392 | 4.262424993 | 0.4735245 | 0.515819892 | 0.491413203 | 29 | 21.47788139 | 20.50715269 | 20.88943586 | 34.87005252 | 19.77589792 | 44.7515153 |
| 11TSL190 | -31.47392413 | 7.233624158 | 4.500095313 | 0.5921914 | 0.648232928 | 0.553106768 | 25 | 22.81247666 | 24.50300682 | 18.02956066 | 38.09570164 | 21.20420254 | 44.79829151 |
| 11TSL191 | -32.14498563 | 7.100596548 | 4.910784634 | 0.49300745 | 0.516966691 | 0.460468958 | 31 | 23.15868747 | 26.41949731 | 18.68400504 | 34.46117152 | 21.29189865 | 42.8387329 |
| 11TSL192 | -31.81798496 | 6.935414487 | 4.736855082 | 0.507437 | 0.508343138 | 0.473946158 | 27 | 22.92618363 | 25.77919782 | 19.01228662 | 33.70774515 | 22.45478044 | 47.62483084 |
| 11TSL193 | -31.33197958 | 6.814043327 | 4.262424579 | 0.4698447 | 0.497326182 | 0.43883495 | 27 | 21.97597948 | 27.26478219 | 18.02931024 | 35.74735325 | 21.58463596 | 44.5611209 |

| Code | Postorbital length (PSL)* | Orbital length (OOL)* | Preorbital length (POL)* | Maxillare length (MXL)* | Lower jaw length (LJL)* | Maxillare width (MXW)* | Pectoral fin length (PCL)* | Pelvic fin length (PVL)* | Dorsal fin height (DOH)* | Head depth (HDD)* | Body depth (BDD)* | Caudal peduncle depth (CPD)* |
|---|---|---|---|---|---|---|---|---|---|---|---|---|
| 11TSL100 | 16.783693 | 8.262831032 | 16.06463522 | 9.749992039 | 8.41175834 | 2.636222477 | 30.1676968 | 25.85111745 | 30.03929506 | 16.88014101 | 38.75028852 | 12.77975199 |
| 11TSL101 | 18.96468112 | 9.499101926 | 16.51352657 | 10.46254204 | 8.645996022 | 3.970625751 | 26.59680532 | 22.79894644 | 28.22493131 | 17.33222284 | 39.53307744 | 12.89821836 |
| 11TSL102 | 19.41602411 | 8.679280621 | 16.20757023 | 10.27038153 | 8.318295072 | 3.524134146 | 27.4514612 | 20.66549323 | 24.23752177 | 18.26694716 | 40.70194006 | 13.57073771 |
| 11TSL103 | 21.09807508 | 10.92179726 | 17.36083562 | 12.18831333 | 10.81198376 | 4.150997702 | 25.33650167 | 21.64228652 | 21.81132791 | 19.66452672 | 39.87927561 | 12.29169421 |
| 11TSL104 | 19.3594268 | 9.071522345 | 15.8937696 | 9.912587978 | 8.128377928 | 4.174211968 | 27.83789257 | 24.88699184 | 29.02471845 | 17.7236869 | 35.02889597 | 11.77668644 |
| 11TSL105 | 20.15186884 | 9.198417049 | 16.01367383 | 10.70230891 | 9.698820919 | 3.566341073 | 27.60719057 | 20.02672356 | 23.60427495 | 18.67475924 | 41.12195958 | 12.90039237 |
| 11TSL112 | 19.05292953 | 10.61358519 | 17.88754931 | 10.27776161 | 7.938194117 | 3.728233 | 28.55021076 | 25.41612083 | 28.21008594 | 17.42153153 | 36.92189294 | 12.614773 |
| 11TSL113 | 20.45060372 | 9.981388632 | 18.2548948 | 11.3072008 | 8.28858736 | 3.522769801 | 29.68404943 | 26.45637523 | 33.05559758 | 18.50841194 | 36.16178605 | 13.30406911 |
| 11TSL114 | 21.13151656 | 11.14168117 | 16.13072974 | 11.37022122 | 7.671372688 | 3.480089816 | 29.45644833 | 26.70593194 | 30.72902363 | 18.65527414 | 37.51093148 | 13.54713331 |
| 11TSL117 | 20.01693363 | 9.717777607 | 16.30198269 | 10.30991832 | 7.313295875 | 3.182845043 | 25.3481528 | 21.9239344 | 27.10167519 | 17.6250989 | 36.50929967 | 13.26553386 |
| 11TSL120 | 20.08100549 | 11.03206208 | 14.33339925 | 11.28208565 | 7.443902975 | 3.159420247 | 26.93088364 | 25.49633249 | 29.41366988 | 17.99963198 | 36.003988 | 12.4102137 |
| 11TSL121 | 20.15110384 | 9.958640993 | 16.43521167 | 10.61831734 | 8.014799345 | 3.47138679 | 23.11905463 | 22.98401431 | 26.06669352 | 17.5687172 | 36.09014322 | 12.63551075 |
| 11TSL122 | 19.05173831 | 11.55743994 | 20.97819858 | 11.6072465 | 8.752234212 | 3.449340682 | 20.6351962 | 23.86231141 | 18.11427504 | 19.14350752 | 41.16894535 | 13.22877456 |
| 11TSL123 | 18.66773873 | 9.210920877 | 15.91256945 | 13.37167439 | 9.801753016 | 4.067137249 | 25.83201647 | 24.08459669 | 27.60691749 | 17.83455726 | 39.08315706 | 13.30928923 |
| 11TSL124 | 18.27936133 | 9.345228818 | 18.95791894 | 9.278574498 | 8.822199766 | 3.951089921 | 28.93313552 | 24.77599551 | 30.67526603 | 16.55895585 | 35.43904064 | 12.5289276 |
| 11TSL127 | 19.1961739 | 9.524606168 | 16.66678292 | 8.62412191 | 8.328530912 | 3.920287129 | 28.80129096 | 24.19453465 | 28.60365359 | 17.82334366 | 39.84563304 | 12.51592619 |
| 11TSL128 | 19.45888087 | 11.86035043 | 16.17064708 | 9.585377293 | 6.6543053 | 3.769402339 | 29.13257722 | 25.13702069 | 31.14832886 | 17.50923586 | 37.76443305 | 13.58560515 |
| 11TSL133 | 20.13772814 | 10.5970461 | 18.39016857 | 9.977325649 | 8.326748479 | 4.348835375 | 29.3901699 | 24.28456792 | 28.02756738 | 18.46289153 | 37.12739364 | 12.57582142 |
| 11TSL134 | 17.94230894 | 9.346311955 | 17.28593521 | 9.495462661 | 7.010504642 | 3.506637966 | 20.16961972 | 21.64269155 | 30.18407971 | 15.9853878 | 34.24644514 | 11.24962487 |
| 11TSL135 | 18.73350745 | 9.167878738 | 21.08652149 | 10.70550392 | 8.407468221 | 3.879181291 | 31.28739232 | 27.06564467 | 31.6140074 | 18.88966076 | 40.06401329 | 13.20757936 |
| 11TSL136 | 21.7085658 | 9.107508952 | 21.69297097 | 11.53563876 | 8.899374713 | 4.025740576 | 28.16562676 | 26.51917662 | 28.92178463 | 17.96846236 | 35.39369916 | 12.8809072 |
| 11TSL137 | 21.27252373 | 8.849234143 | 15.13349808 | 9.680966342 | 8.052332164 | 3.677663247 | 31.05172473 | 23.15669436 | 29.81399302 | 17.60646175 | 36.38642753 | 12.22338239 |
| 11TSL138 | 20.22219396 | 10.09758394 | 18.46022295 | 10.15300799 | 8.29235098 | 4.079150316 | 28.06231813 | 23.65422277 | 27.99674372 | 17.68043196 | 34.76332171 | 11.95510919 |
| 11TSL139 | 19.24643922 | 8.932279926 | 14.89675823 | 8.856172597 | 6.330173388 | 3.40798653 | 26.70507288 | 23.78493452 | 26.62781197 | 17.02501619 | 39.66299128 | 12.71208715 |
| 11TSL140 | 22.18397523 | 9.880738344 | 18.50743777 | 11.83834475 | 9.533506639 | 3.931477318 | 33.24216599 | 27.80032921 | 31.78799932 | 19.64183937 | 38.91756533 | 13.69598709 |
| 11TSL146 | 21.40036128 | 9.450965814 | 21.05879854 | 10.3128851 | 8.309254827 | 3.893872601 | 28.08780363 | 22.8074392 | 29.29711859 | 17.59770809 | 39.92689605 | 13.45405396 |
| 11TSL147 | 20.61109306 | 10.14824158 | 17.59294286 | 9.397742618 | 7.202880931 | 3.410088742 | 29.76554245 | 25.57093019 | 30.89031732 | 18.26372145 | 40.06899977 | 12.54472825 |
| 11TSL148 | 19.97369772 | 9.57014323 | 18.20290405 | 8.871628966 | 7.113440262 | 4.165200561 | 30.12598849 | 23.49996679 | 28.47191688 | 18.12245899 | 37.7870668 | 12.66650478 |
| 11TSL150 | 20.61188252 | 9.42840656 | 18.21438125 | 10.61279363 | 8.844625563 | 3.288847166 | 28.76955233 | 26.62698935 | 33.49227066 | 18.00076219 | 40.40798662 | 12.67103519 |
| 11TSL151 | 16.70361921 | 9.097956729 | 15.80482056 | 8.667233467 | 6.283096843 | 3.465956296 | 29.77633331 | 24.15151683 | 30.4431457 | 15.88087097 | 37.16007323 | 11.99103197 |
| 11TSL152 | 18.96455536 | 8.893201598 | 18.14869955 | 9.742353114 | 8.146292323 | 3.770817884 | 28.34171844 | 26.71218246 | 31.74423567 | 16.08221986 | 37.22840101 | 12.63414076 |
| 11TSL153 | 19.89188996 | 9.65517166 | 18.68320416 | 9.964952212 | 8.888845476 | 3.974248993 | 30.11129054 | 25.94092432 | 30.10786313 | 18.34826714 | 39.2006943 | 13.21108906 |
| 11TSL154 | 20.16492767 | 8.294833591 | 21.90354191 | 9.065204846 | 7.032511097 | 4.129883118 | 28.7973504 | 23.20055828 | 30.94949162 | 16.9204593 | 36.8042751 | 12.41989686 |
| 11TSL155 | 21.08625545 | 9.083780054 | 17.72781432 | 11.06921913 | 8.922886723 | 4.102396473 | 26.60172535 | 28.28383869 | 32.40259703 | 19.1496055 | 40.75540025 | 12.60170181 |
| 11TSL156 | 20.21194189 | 9.715716173 | 18.06144769 | 10.72203566 | 8.79684855 | 3.646667909 | 32.12721232 | 26.14830241 | 31.97467269 | 17.8425313 | 37.76983334 | 13.12697641 |
| 11TSL157 | 18.42263013 | 10.25507908 | 22.54028929 | 11.10635555 | 8.615821134 | 3.407373007 | 29.81301253 | 24.23937886 | 28.67990347 | 18.19154665 | 37.36562485 | 12.00839685 |
| 11TSL159 | 20.2139819 | 9.81151919 | 18.44797601 | 10.42971471 | 8.098609201 | 4.19613343 | 30.10587854 | 25.2889638 | 30.46581684 | 18.23559995 | 34.6868426 | 12.02336007 |
| 11TSL160 | 19.68035661 | 9.178984193 | 16.61862526 | 9.582981826 | 8.109363213 | 4.30023679 | 29.76236404 | 26.49923382 | 31.52288834 | 18.51321266 | 38.39370645 | 12.91226897 |
| 11TSL161 | 20.95711983 | 9.090242377 | 22.94199625 | 9.772650983 | 9.838495466 | 3.256677778 | 30.57618009 | 25.59658056 | 30.2222336 | 17.11274028 | 39.47682416 | 12.87563375 |
| 11TSL170 | 18.94154789 | 10.54604066 | 18.25766437 | 9.441909309 | 7.121182443 | 4.019331092 | 31.21213646 | 25.42900074 | 29.42129304 | 19.53458827 | 37.59665259 | 13.11243804 |
| 11TSL171 | 19.13055943 | 9.364647462 | 19.03298529 | 9.39364164 | 7.32083926 | 3.986639804 | 28.72981328 | 24.97838582 | 27.34432682 | 16.20526089 | 36.13638479 | 11.62448264 |
| 11TSL172 | 20.07406992 | 9.718325348 | 16.78005965 | 10.27713701 | 8.538088992 | 3.836155316 | 30.43893759 | 25.88662377 | 30.37513362 | 16.51351913 | 38.79370387 | 11.80191396 |
| 11TSL173 | 18.72567631 | 10.58529662 | 16.49820797 | 10.55479453 | 8.21648519 | 3.702883558 | 26.08232595 | 22.69818778 | 28.54456141 | 16.45955505 | 39.01994573 | 12.38376658 |
| 11TSL174 | 19.11176083 | 10.83454677 | 13.84823152 | 11.171111 | 8.942672592 | 4.186015774 | 27.3283035 | 24.30527122 | 27.51670911 | 17.60051708 | 39.78429097 | 12.39935899 |
| 11TSL176 | 19.87668052 | 8.916802772 | 16.62892555 | 8.288749851 | 6.801545388 | 3.491879795 | 30.94942289 | 25.27829806 | 33.85888466 | 16.53446294 | 41.97011298 | 13.23154102 |
| 11TSL178 | 21.25615919 | 9.542891037 | 15.52426438 | 10.16567771 | 7.706019795 | 4.507870665 | 33.56459969 | 28.53348914 | 31.52545921 | 16.91924012 | 38.567831 | 13.02868024 |
| 11TSL179 | 20.50342616 | 9.453437901 | 19.3403944 | 10.42815282 | 7.744023855 | 3.45300028 | 30.85470711 | 25.52601067 | 29.05521174 | 18.42575772 | 44.15762919 | 13.42217349 |
| 11TSL180 | 18.43513136 | 10.28393903 | 17.20406974 | 10.87800932 | 8.412274537 | 4.223108689 | 29.67194932 | 25.51413559 | 29.52952999 | 17.56631052 | 38.78527222 | 12.57010343 |
| 11TSL181 | 20.32133915 | 9.96015217 | 17.36157468 | 10.84190263 | 8.884615962 | 4.071106232 | 31.92366702 | 25.44717677 | 30.12403485 | 17.8416581 | 39.24262266 | 12.98404971 |
| 11TSL182 | 21.69520316 | 9.822681536 | 17.59299171 | 9.366568376 | 6.888072164 | 4.207950601 | 29.85661251 | 23.1239439 | 28.40012886 | 17.76652766 | 39.22641735 | 12.02073215 |
| 11TSL183 | 19.10393174 | 9.155435437 | 16.46321058 | 9.965604451 | 8.004058107 | 3.399583554 | 29.03614087 | 24.9078259 | 28.39786201 | 16.74937396 | 39.80733912 | 12.68245537 |
| 11TSL184 | 19.26354973 | 10.69864645 | 16.50257999 | 9.679180909 | 7.318134078 | 4.240950981 | 29.37747474 | 26.41405682 | 30.59304359 | 17.2758871 | 44.23644756 | 12.44252647 |
| 11TSL185 | 20.06075696 | 10.00802772 | 17.01487344 | 9.410547832 | 7.874355685 | 3.861255395 | 27.7003227 | 25.28194211 | 29.37585486 | 16.49613489 | 36.67864444 | 12.32921816 |
| 11TSL186 | 20.33613733 | 9.002849423 | 17.40796173 | 11.14824584 | 9.236038787 | 4.061372619 | 30.48154287 | 25.97631417 | 31.37062902 | 17.34337249 | 39.37980948 | 13.21913802 |
| 11TSL187 | 20.44958409 | 9.799961331 | 19.50311493 | 10.1230983 | 8.36644235 | 3.819607592 | 29.24507172 | 24.6233783 | 29.40413166 | 17.19651076 | 40.19667913 | 13.19619908 |
| 11TSL188 | 18.70378422 | 9.650563318 | 19.62805653 | 9.646595546 | 8.660947344 | 3.922473551 | 28.01468003 | 24.16878648 | 28.72190979 | 17.22326342 | 39.41720262 | 12.17701509 |
| 11TSL189 | 18.57495996 | 9.313955093 | 18.3390963 | 8.962967424 | 6.374187342 | 3.925106025 | 28.99778169 | 25.39309286 | 30.26701404 | 15.94273256 | 37.88780917 | 13.08412353 |
| 11TSL190 | 19.45994546 | 9.766261198 | 17.79832808 | 10.42074718 | 7.407603753 | 3.310970541 | 27.23358427 | 23.71546526 | 26.67024232 | 17.02428192 | 36.21559766 | 12.75934226 |
| 11TSL191 | 19.92334057 | 10.42057238 | 17.55226037 | 9.467476716 | 7.661562647 | 3.827474529 | 28.32204382 | 23.34277309 | 27.94338904 | 17.6077425 | 36.79448371 | 11.81692091 |
| 11TSL192 | 19.81427571 | 10.05129171 | 17.29252791 | 9.165078925 | 7.627951213 | 3.681544803 | 29.44320204 | 24.62932674 | 29.86777028 | 17.33941035 | 38.76477415 | 12.7274444 |
| 11TSL193 | 18.39724955 | 9.60244232 | 15.36708067 | 10.7328738 | 8.777627375 | 3.752728239 | 30.98843785 | 26.02770176 | 30.22892181 | 17.45790807 | 39.34998445 | 13.12960978 |

***total length adjusted**

****age adjusted**

**Table S2. Summary of scaled demographic estimates based on the best-fit model (SC_NeC) through LRT tests.** The Modelled 2D-SFS based on SC_NeC is given in Figure S4. Abbreviations of demographic parameters are expanded in Figure S5.

| Parameters | Estimate | 95% C.I. | S.D. |
|---|---|---|---|
| **theta** | 2401.23243 | 2277.18562–2525.27923 | 488.4647 |
| **nu1** | 4.06934 | 1.62684–6.51183 | 9.61792 |
| **nu2** | 1.80447 | 1.56178–2.04716 | 0.95564 |
| **nu1b** | 0.57837 | 0.51154–0.6452 | 0.26315 |
| **nu2b** | 0.38192 | 0.33115–0.43269 | 0.19993 |
| **T1** | 1.81654 | 1.59799–2.0351 | 0.86061 |
| **T2** | 0.12569 | 0.10876–0.14262 | 0.06666 |
| **m12** | 14.24471 | 13.14531–15.34412 | 4.32919 |
| **m21** | 9.59325 | 7.85582–11.33067 | 6.84154 |
| **O** | 0.93697 | 0.93387–0.94008 | 0.01223 |
| **Nref** | 16633.30233 | 15774.03189–17492.57277 | 3383.58792 |
| **T1Gen** | 55234.81209 | 52768.22862–57701.39555 | 9712.77683 |
| **T2Gen** | 3995.2187 | 3488.83862–4501.59878 | 1993.99564 |
| **T1Year** | 276174.0604 | 263841.14312–288506.97776 | 48563.88413 |
| **T2Year** | 19976.0935 | 17444.19311–22507.99389 | 9969.97822 |
| **T12_ratio** | 16.09971 | 14.64694–17.55247 | 5.72062 |
| **Ttot** | 1.94224 | 1.71544–2.16903 | 0.89305 |
| **TtotGen** | 59230.03079 | 56891.4909–61568.57068 | 9208.57389 |
| **TtotYear** | 296150.1539 | 284457.45448–307842.8534 | 46042.86944 |
| **N1** | 74463.89416 | 25548.92383–123378.86449 | 192614.6823 |
| **N2** | 28001.33201 | 24671.86872–31330.7953 | 13110.57758 |
| **N1b** | 8980.41332 | 8518.94838–9441.87826 | 1817.13127 |
| **N2b** | 5845.47176 | 5508.55405–6182.38947 | 1326.69604 |
| **N1end** | 8980.41332 | 8518.94838–9441.87826 | 1817.13127 |
| **N2end** | 5845.47176 | 5508.55405–6182.38947 | 1326.69604 |
| **M12** | 0.00043 | 4e–04–0.00046 | 0.00012 |
| **M21** | 0.00027 | 0.00024–0.00031 | 0.00013 |
| **Nem12** | 7.08661 | 5.3677–8.80551 | 6.76861 |
| **Nem21** | 21.29215 | 10.95441–31.62988 | 40.70736 |

**Method S1.**

*Fish sampling and measurements*

Fish were sampled with gill net series, comprised of eight 1.8 m high and 30 m long gill nets (knot to knot mesh size 12, 15, 20, 25, 30, 35, 45, 60 mm) and one multimesh Nordic gill net (height 1.5 m, length 30 m, mesh sizes 5-55 mm), to capture different sized fish. Gill net series was set in the evening to benthic zone in littoral (depth < 6 m), benthic zone in profundal (depth > 10 m) or open-water pelagic 0-4 m depth with floats. Soaking time was circa 12 hours and fish were immediately killed during the gill net lifting and ice was applied to cooling down the fish.

Fish were removed from gill nets and sorted to habitat and species. Capture habitat (littoral, pelagic, profundal), total length (accuracy of 1 mm) and weight (0.1 g) were measured from each whitefish individual and samples were frozen to -20°C for subsequent analyses. Fish were later melted and pinned for 17 body morphometric measurements (accuracy 0.001 mm) (see Figure S2). After linear body measurements, the first left gill arch was dissected for gill raker counting, gill arch and longest gill raker measurements under preparation microscope (Kahilainen & Østbye 2006). Body morphometric and gill raker measurements were corrected to mean total length of 22.4 cm of all individuals (Kahilainen *et al.* 2011).

Sagittal otoliths and scales between ventral and anal fins were taken for age determination. Dorsal white muscle tissue and liver sample was stored to plastic vial for later analyses. Sex and sexual maturity were visually determined and both gonads were weighed (accuracy 0.01 g). Gonads were adjusted to mean total length of all whitefish samples. Stomach content was studied with points method (Hynes 1950) at scale 0-10, where 0 defines empty stomach and 10 is extended full stomach. The proportion of each prey category to total stomach fullness was visually determined. Diet was defined as pelagic (sum of *Bosmina* sp., Chironomidae pupae, terrestrial insects, Copepoda, Calanoida) or benthic (sum of Trichoptera larvae, *Valvata* sp., Chironomidae larvae, *Lymnaea* sp., *Pisidium* sp., *Eurycercus* sp., Ostracoda, Hydracarina) to calculate the proportion of pelagic prey. Pelagic parasite abundance, i.e. cysts *Diphyllobothrium* spp., was counted from stomach wall following Kahilainen *et al.* (2011).

Dorsal muscle and liver tissues were freeze-dried for 48 hours at -70°C and later ground for fine powder. Subsequently, circa 1 mg of tissue powder was weighed and encapsulated to a tin cup for stable isotope analyses of carbon ($d^{13}C$) and nitrogen ($d^{15}N$) was run with constant flow elemental analyser (PDZ Europa ANCA-GSL) connected with stable isotope ratio mass-spectrometer (PDZ Europa 20-20). Instrumental error for both carbon and nitrogen was 0.2 ‰. Stable isotope ratio was given as raw values and no lipid correction was conducted. Lower $d^{13}C$ values <-27‰ indicates pelagic/profundal resource use and higher values >-27 ‰ typically indicates littoral habitat use. Stable isotope of nitrogen indicates trophic level that is typically lower in littoral invertivorous fish feeding on shorter food chains than pelagic planktivores (Harrod *et al.* 2010). Profundal benthivores tend to have high $d^{15}N$ ratio due to their reliance on setting material from pelagic (Harrod et al.

2010). Elemental ratio of carbon and nitrogen was compared as increasing value indicates more lipids (see Kahilainen *et al.* 2017).

Total mercury was run from freeze-dried dorsal muscle powder (20-30 mg per sample) using direct mercury analyser (DMA-80, Milestone, Italy). Each sample was run as duplicate and the mean value of duplicate was used in subsequent analyses. Instrumental error was assessed by using blank sample vial and certified reference material (DORM-4, Canadian Research Council) at the beginning and end of each run (see Kahilainen *et al.* 2017). In subarctic lakes, littoral invertivorous whitefish tend to have low mercury content due to low mercury in their prey and short food chains, whereas pelagic and profundal invertivorous whitefish tend to have higher mercury content (Kahilainen *et al.* 2017). As mercury accumulate to fish with increasing size and age, we corrected the values to mean total length (22.4 cm) and age (9.3 years) (see Kahilainen *et al.* 2017).

*Genome-wide genetic divergence*

We calculated unfolded two-dimensional site frequency spectrum (2D-SFS) based on the sites shared between pairwise ecotypes with ANGSD. The $F_{ST}$ statistics for pairwise ecotypes was calculated through function "fs.Fst" in *Moments* (Gutenkunst *et al.* 2009) and the $d_{XY}$ statistics was calculated using a custom R script from Momigliano et al. (2021; Zenodo data repository: doi: 10.5281/zenodo.451837). To calculate $D_a$ ($D_a = d_{XY}$ -[$\pi_{\text{population1}}$-$\pi_{\text{population2}}$]/2, Nei (1987); $\pi$, nucleotide diversity, Nei & Li (1979)), we firstly marginalized 2D-SFS to 1D-SFS per population using function "fs.marginalize", followed by calculating $\pi$ per population with the function "fs.pi" in *Moments*.

*Demographic modelling*

The demographic history of the littoral and pelagic ecotypes (profundal ecotype was not analysed as it was not found to be significantly divergent from the pelagic ecotype; see Methods) was reconstructed in *Moments* based on the 10 simulated two-population models (Figures S5) using unfolded two-dimensional SFS (2D-SFS). All models were optimized in 10 optimization routines, each consisting of five rounds of optimization using the approach from Momigliano *et al.* (2021)) similarly to Portik *et al.* (2017)). Among the 10 models, we tested and chose the best model based on Likelihood Ratio Test (LRT) by adjusting the D-statistics to account for possible effects of linkage (Coffman *et al.* 2016). To find the best model, we firstly tested whether the extended model outperforms the basic one for each demographic scenario, such as SI *vs.* SI_NeC, and then compared all different models reciprocally, for example SI_NeC *vs.* AM_NeC. Based on the best model (SC_NeC in this study), we eventually inferred the 95% confidence interval (CI) of the demographic parameters by bootstrapping the input 2D-SFS 100 times with ANGSD, permuting loci within chromosome (realSFS -bootstrap 100 -resample_chr 1). Parameters in coalescent units were scaled based on the simulated mutation rate $\mu$=1×10$^{-8}$ and an assumed generation time of 5 years for whitefish (Thibert-Plante *et al.* 2020) using methods outlined in *Moment's* manual. The scripts for specifying models, *Moments* analyses and LRT tests were deposited in Zenodo repository (Data accessibility).

*Admixture mapping*

To associate SNPs with trait variation, we conducted GEMMA and LDna-EMMAX analyses for the 41 admixed individuals. For GEMMA, a VCF file was produced using ANGSD with the following filters: -minMaf 0.05, -minInd 30, -postCutoff 0.95 -doVcf 1, -SNP_pval 1e-6. Missing data were imputed using BEAGLE (Browning & Browning 2007) and a binary Plink BED format was generated using VCFtools (Danecek *et al.* 2011). A relatedness matrix was produced and used as a covariate in GEMMA (-gk). We performed univariate linear mixed models for each trait in GEMMA and adopted a significance threshold of $\alpha = 0.00001$ ($-\log_{10}(\alpha) = 5$) in Wald test (-lm 1).

LDna-EMMAX was conducted following the pipelines in Kemppainen *et al.* (2021)). Specifically, we started grouping loci connected by high linkage disequilibrium (LD) into LD-clusters using LD network analyses (LDna; Kemppainen *et al.* (2015)) based on genotype likelihoods. Next, the principal component analysis (PCA) was performed for each LD-cluster using *PCAngsd*, followed by linear mixed models (LMMs) testing for associations between the first PC (PC1) of the LD-clusters and traits. The association analyses were corrected for *P*-value inflation by including relatedness as a random effect (Li *et al.* 2018; Fang *et al.* 2021; Kemppainen *et al.* 2021) and for multiple testing through permutation (Li *et al.* 2018). R-scripts of LDna-EMMAX used in this study were available in Zenodo repository (see Data accessibility statement).
